# Regionally enriched rare deleterious exonic variants in the UK and Ireland

**DOI:** 10.1101/2022.09.19.508526

**Authors:** Mihail Halachev, Elvina Gountouna, Alison Meynert, Regeneron Genetics Center, Gannie Tzoneva, Alan R. Shuldiner, Colin A. Semple, James F. Wilson

**Affiliations:** MRC Human Genetics Unit, Institute of Genetics and Cancer, University of Edinburgh, United Kingdom; Regeneron Genetics Center, Tarrytown, NY, USA; Centre for Global Health Research, Usher Institute, University of Edinburgh, United Kingdom

## Abstract

Geographic clustering of haplotypes appears to have emerged in the UK as a result of differing patterns of immigration and drift in regions that have been relatively isolated from each other. However, until recently it has been unclear how such patterns of regional genetic differentiation might impact the protein-coding fraction of the genome. Here, we exploit UK Biobank (UKB) and Viking Genes whole exome sequencing data to study regional genetic differentiation across the UK and Ireland in protein coding genes, encompassing 20 regions of origin and 44,696 unrelated individuals. We rediscover the strong influence of genetic drift in shaping variation in the Northern Isles of Scotland and among those with full or partial Ashkenazi Jewish (AJ) ancestry. For full AJ, almost half the known rare exonic variants (45%) are at least two-fold more or less frequent than in a Europe-wide reference sample, while the degree of variant frequency differences in Shetland and Orkney are comparable to part AJ (19%, 17%, 16%, respectively). We also demonstrate substantial genetic differentiation among several mainland regions of origin, particularly north and south Wales, SE Scotland and Ireland. With stringent filtering criteria we found 67 variants likely to have adverse biomedical consequences, enriched by at least five-fold in frequency in one or more British or Irish regions relative to a European reference group, and we calculate that this may lead to tens or hundreds of affected individuals. We conclude that regional genetic variation across the UK and Ireland should be considered in the design of genetic studies, and may inform effective genetic screening and counselling.

## Introduction

Geographically diverse human populations often exhibit distinct profiles of genomic variation. This was first established based on mitochondrial DNA^1^ and Y chromosome haplotyping^2,3^ and the advent and mass adoption of next-generation sequencing technologies soon made clearer the true breadth and complexity of this phenomenon. For example, the 1000 Genomes Project conducted whole-genome sequencing (WGS) and analysis of 2504 individuals from 26 populations in Africa, East Asia, Europe, South Asia and the Americas and found that rarer variants are typically restricted to closely related populations, with the vast majority of variants found only in a single continental group^4^. Such differential signal persists even at smaller geographical distances and is evident even when only a subset of the full genomic variation is investigated. Analysing genome-wide single nucleotide polymorphism (SNP) genotyping data of 2039 individuals from rural areas within the UK (and with grandparents within the same areas), Leslie et al.^5^ showed remarkable concordance between genetic and geographic clustering of samples across the country. Further differentiation was reported by Gilbert at al.^6^ based on genome-wide SNP genotyping data analysis of 2544 individuals from five different cohorts of regional English, Welsh, Scottish, Manx, or Irish ancestry.

Isolated populations can show more extreme divergence due to strong genetic drift. The European Ashkenazi Jewish (AJ) population has long been regarded as a genetic isolate showing clear evidence for genetic drift arising from population bottlenecks, endogamy, as well as complex patterns of admixture and selection at particular loci^7^. We have previously found strong genetic drift in the isolated Shetland population in northern Scotland, relative to the more cosmopolitan mainland Scottish population^8^. Many of the ultra-rare exonic variants found to be enriched in Shetland are predicted to impact gene function and may affect biomedical traits^9^, consistent with similar enrichments observed in other geographically isolated populations^10–13^. The Shetland population’s demographic history reflects the substantial physical barriers to immigration historically, and it is thought that over a similarly recent timescale many regions of the UK may have experienced limited migration^14,15^, preserving regional genetic clusters that appear to reflect more ancient histories of those regions^5,16^. However, it is unclear whether these patterns of regional differentiation have any relevance to health and disease.

UK Biobank (UKB) is a large-scale biomedical database providing medical and genetic data to accredited researchers from half a million volunteer participants with the aim of enabling new scientific discoveries and improving public health^17^. A UMAP analysis of the genome-wide SNP data of 488,377 UKB participants confirmed the previously observed regional genetic stratification in the UKB, showing clear clustering based on self-reported ethnic background, as well as north-south and east-west gradients^18^. This large-scale and richly annotated dataset has already provided valuable insights into human health and disease. A plethora of association studies revealed numerous genotype-phenotype associations for common/complex diseases, with some focusing on the effect of common variants^19–21^, while others investigated the contribution of rare variation^22,23^.

In this work, using a subset of the whole-exome sequencing (WES) data available for UKB participants^24^ focused on self-identified “White British” individuals born outside large metropolitan areas, along with that from Viking Genes participants from the Northern Isles of Scotland, we sought to address three main questions. Is the genome-wide signal driving the previously observed population differentiation in the UKB also seen in the protein-coding fraction of the genome? Are distinct spectra of rare exonic variants observed in the genetic clusters associated with different regions? Are there regionally enriched exonic variants of biomedical importance and what are their practical implications?

## Results

Based on the regional availability of participants with WES data in UKB we classified samples into 16 geographical regions of origin (Methods). These regions contain individuals who were born within the corresponding region, but outside large metropolitan areas, who self-identify as “White British” and who exhibit very similar genetic ancestry based on a principal components analysis of the UKB whole-genome SNP array genotypes. There are two exceptions: the London region which contains individuals born in a 10 mile radius area around the geographical centre of London (i.e., a cosmopolitan control) and the Irish region for which we selected individuals who self-identify as “Irish” and were born in either Northern Ireland or the Republic of Ireland. We also included UKB participants with Ashkenazi Jewish (AJ) heritage, which we split into two groups (full and part AJ) based on their genomic information (Methods). Lastly, we added WES data from two cohorts in the Viking Genes programme^9,25^, from the relatively isolated archipelagos of Shetland and Orkney (the Northern Isles of Scotland), for which the sequencing and variant calling procedures were identical to those utilized for UKB WES data generation, for a total of 20 regions and 44,696 unrelated individuals (Fig S1).

### Individuals with Ashkenazi Jewish heritage in UKB

According to the 2011 UK census, more than quarter of a million respondents answered “Jewish” to the voluntary question on religion. Recent studies have found evidence of participation of such individuals in the UKB project, including a study based on identity-by-descent (IBD) analysis of the 500k UKB participants^26^ and a recent analysis of European haplotype sharing in UKB SNP genotyping data^27^. An independent clustering analysis based on UKB whole-genome SNP array genotypes also revealed a distinct group of UK individuals, which based on their genetic data and UKB lifestyle questionnaire answers were believed to be of Jewish ancestry [David W. Clark, personal communication]. Our further analysis of these individuals using the UKB WES data indicated that this group is enriched for some known pathogenic variants causing disorders with higher prevalence in Ashkenazi Jewish (AJ) individuals, including a frameshift variant in the *HEXA* gene causing Tay-Sachs disease (rs387906309, ∼50x enrichment in the AJ group compared to Central London) and a missense variant in the *GBA* gene causing Gaucher disease, Type I (rs76763715, ∼13x enrichment). Our WES-based Multi-Dimensional Scaling (MDS) analysis revealed the existence of two main clusters within this group (Fig S1A), which we speculated to be a representation of the varying degrees of admixture between AJ and the remainder of the UK population and which we refer to as full AJ (presumably with 3 or more AJ grandparents) and part AJ (e.g. 2 or less AJ grandparents). Our hypothesis that full AJ individuals have stronger AJ heritage compared to part AJ is supported by two lines of evidence: *a*) an MDS analysis based on known biallelic SNPs which positions the part AJ group between the full AJ and London individuals (Fig S1B); *b*) a smaller total number of runs of homozygosity (ROH)^28^ and a smaller overall proportion of each individual’s genome was observed in ROH for part AJ compared to full AJ (Fig S1C). We included full AJ (1004 unrelated individuals) and part AJ (657 unrelated Individuals) in the following analyses as representative groups of a well-established human isolate population which are at different stages of admixture with other, more numerous groups in the UK population, and serving as archetypal groups enriched for variants that are rare elsewhere.

### Enrichment of shared ultra-rare SNP alleles in the Northern Isles

After performing extensive QC filtering (Methods) of the variants discovered by the UKB alignment and variant calling OQFE protocol in each of the 20 regions, we found that the mainland UKB regions are virtually indistinguishable in terms of overall variation load, with individuals from these regions carrying a median of approximately 31,900 SNP alleles and 823 INDEL alleles in their exomes (Table S1). We annotated variants as “ultra-rare” if they have not been observed in any individual in the full gnomAD genome dataset (v3.1.1, n = 76,156) or “known” if found in any gnomAD subpopulation^29^ with passing variant quality. We further split the known variants based on their frequency in gnomAD Non-Finnish Europeans (NFE, n = 34,029) to “very rare” for variants with MAF_NFE_ < 1%, “rare” with 1% ≤ MAF_NFE_ < 5%, and “common” with MAF_NFE_ ≥ 5%. We note a slight overall enrichment of variants observed in individuals with AJ heritage (∼1% SNP enrichment in full AJ and ∼0.5% in part AJ), mainly driven by an enrichment of very rare known exonic variants, as well as a relative depletion of ultra-rare variants. These variances can be explained by the partial West Asian ancestry in AJ individuals and the inclusion of genomic variation data from 1736 AJ individuals in the gnomAD dataset (2.3% of all 76,156 individuals). We also observed significant enrichment of shared ultra-rare SNP alleles in the Northern Isles, such that two-thirds of the ultra-rare variants found in Shetland are shared by two or more unrelated individuals from this region and more than half of the ultra-rare variants in Orkney are shared among individuals located there. This compares to only one-fifth of ultra-rare SNP alleles observed to be shared among individuals within the London region. This extends previous reports of such enrichments, which have been attributed to founder effects and increased genetic drift in the isolated Shetland population^8^. The effect of the geographical isolation of the Northern Isles is further manifested in the lower number of very rare known SNPs per individual, with its effect potentially underestimated for Orkney, due to the presence of 23 Orcadian individuals in the gnomAD dataset, via their inclusion in the 1000 Genomes project^4^.

### Rare exonic variation is associated with birthplace

Recent research based on genome-wide genotyping arrays has demonstrated a striking association between genomic variation and place of birth for individuals in the UK and the Republic of Ireland^5,6^. To assess if this geographical distinction can be recapitulated based on exonic data only, we assembled a dataset of 10,001 unrelated individuals from the UKB and the Northern Isles (492 Shetlandic, 509 Orcadian and 500 randomly chosen individuals from the remaining 18 groups). Performing MDS/UMAP analyses based upon rare (MAF<0.05) exonic SNP variation (Methods) using the top 20 MDS dimensions reveals a clear distinction of full AJ, part AJ, Shetland and Orkney populations from each other and from mainland regions (Fig 1A, Fig S3A). Focusing on the 16 mainland UK and Ireland regions, distinctions among Welsh, English, Scottish and Irish exomes are evident, consistent with previous studies based on genome-wide genotyping arrays^5,6^ (Fig 1B, Fig S3B). In addition, our analysis reveals an additional differentiation between North and South Welsh individuals, and suggests some level of separation exhibited by individuals born in South East Scotland (Fig 1B). The exonic difference between individuals born in North or South Wales was further confirmed when we restricted the analysis only to Welsh individuals (Fig S4A, Fig S5A); however, performing a mainland Scotland-only analysis did not present convincing evidence that the South East Scotland region was distinct from the other two Scottish regions (Fig S4B, Fig S5B). In contrast, we were not able to find any evidence for exonic distinctions between the individuals from the 10 English regions (Fig S4C, Fig S5C), nor between individuals who self-identify as Irish and were born in either Northern Ireland or the Republic of Ireland (Fig S4D, Fig S5D).

**Figure 1.**
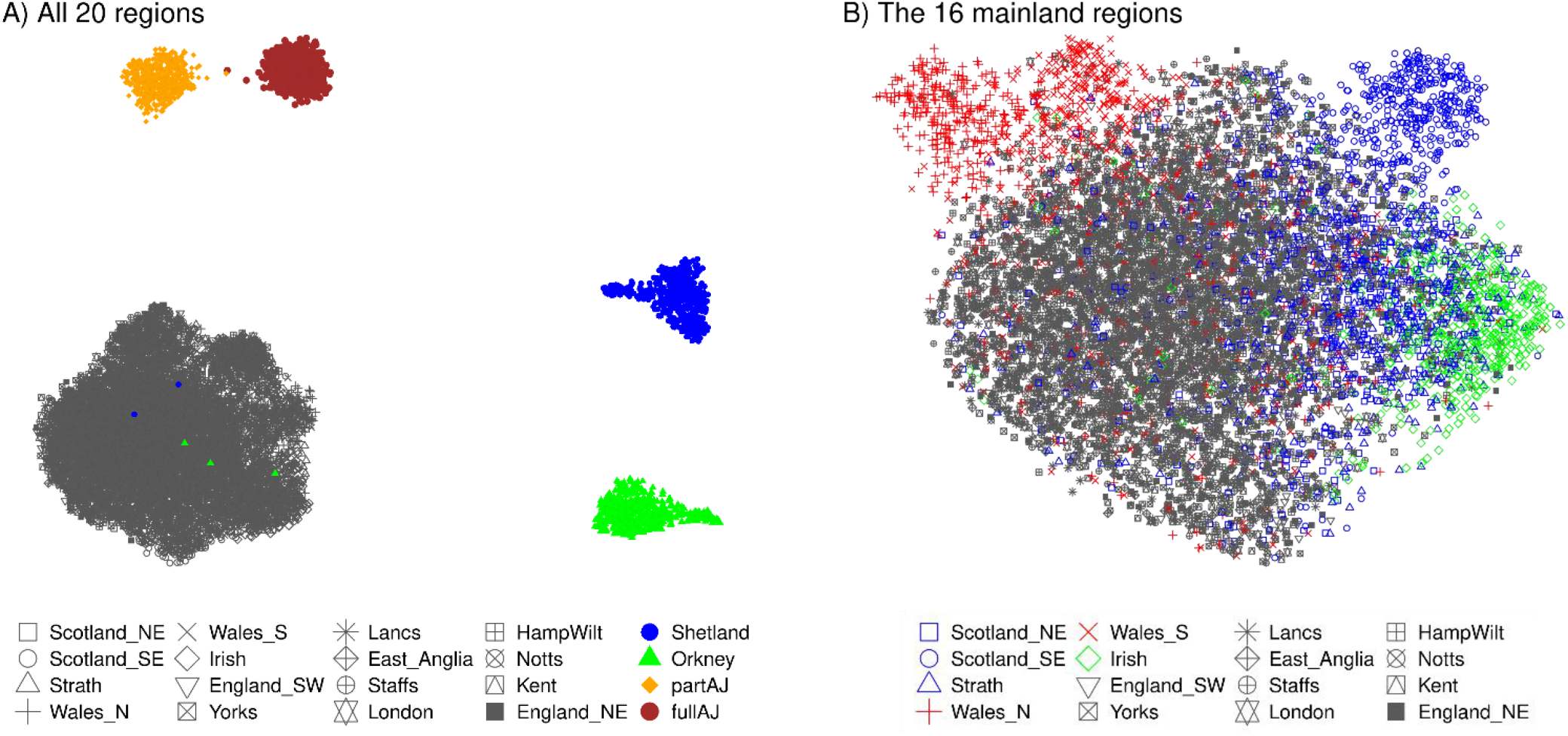
Distinctions among regional populations based upon UMAP projections of rare exonic variation. The UMAP projections are computed on the top 20 MDS dimensions discovered based on biallelic, non-singleton and linkage-disequilibrium (LD) pruned known SNPs with MAF < 5% in the considered unrelated individuals. **A)** UMAP analysis of all 20 groups in our study illustrating the clear genetic distinction of full AJ, part AJ, Shetland and Orkney individuals from each other and from mainland regions. Despite the careful curation of the genealogical records of the Northern Isles participants, some carry a significant proportion of UK mainland heritage; **B)** UMAP analysis focusing on the 16 mainland regions, recapitulating previously known distinctions among Welsh, English, Scottish and Irish regions.

We also computed the pair-wise *F*_*ST*_ distances (Methods) based upon biallelic, non-singleton, linkage-disequilibrium (LD) pruned known SNPs with MAF < 5% across the 20 geographical regions as another measure of the exonic distance between the regions (Table S2). The results further highlighted the clear exonic distinctiveness of the AJ and Northern Isles populations to the 16 mainland regions, suggesting that the individuals from Shetland and Orkney represent a degree of genetic divergence from the mainland regions in their exomes that is comparable (mean *F*_*ST*_ = 0.00090) to the divergence of the part AJ (Fig S6). In accord with the MDS analysis, Irish, Welsh and mainland Scottish regions show elevated mean *F*_*ST*_ distances to each other and to the English (0.00024, 0.00015, 0.00011, respectively), compared to comparisons within England or mainland Scotland (mean *F*_*ST*_ = 0.00005, 0.00004, respectively). The unrooted phylogenetic tree (Fig 2) we built based upon the pair-wise *F*_*ST*_ distances reiterates the Welsh-English-Scottish-Irish differentiation revealed by our MDS/UMAP analysis.

**Figure 2.**
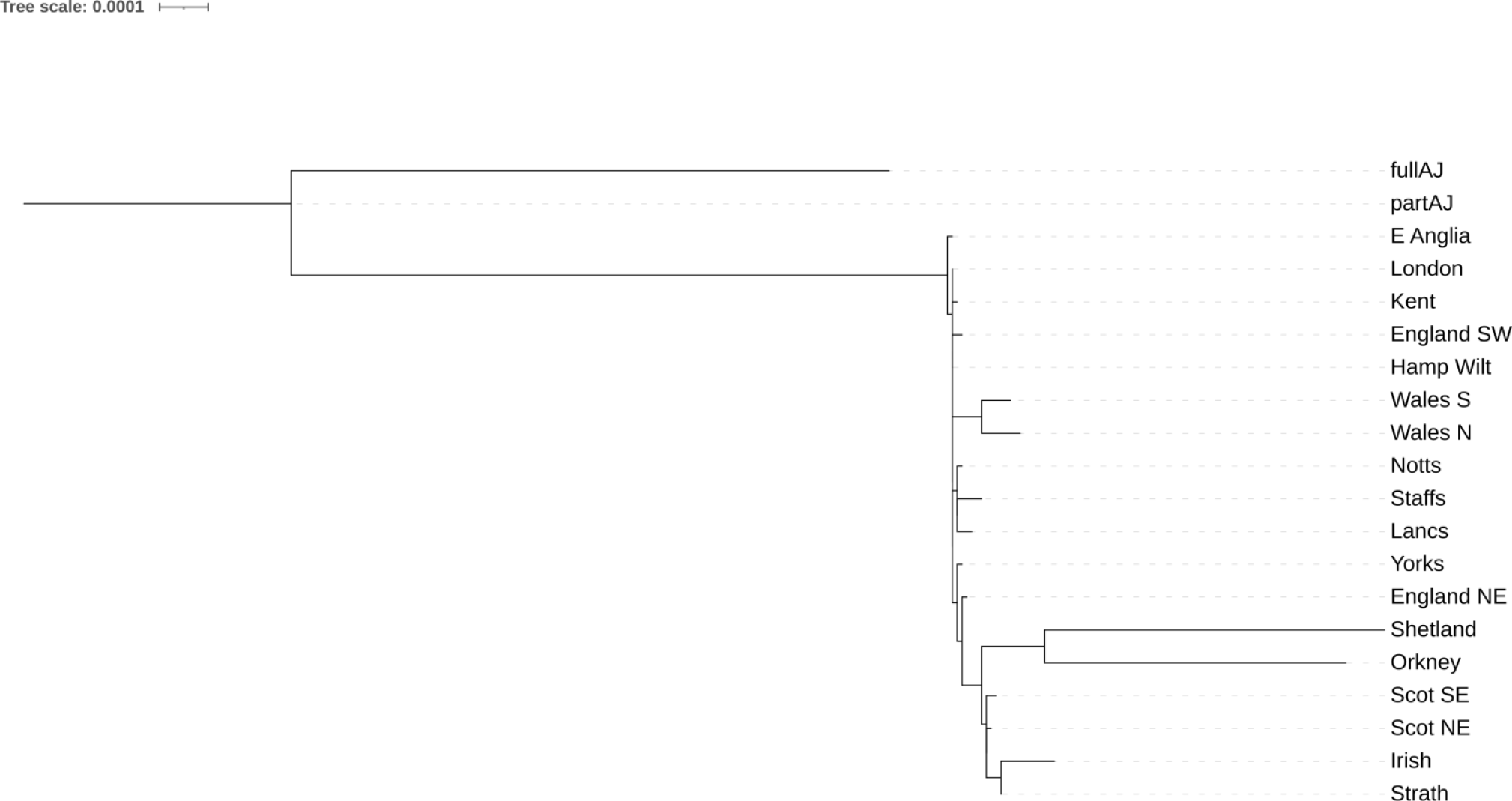
NJ phylogenetic tree for the 20 regions built based upon the pair-wise *F*_*ST*_ distances

### Evidence for genetic drift in mainland regions of the UK and Ireland

Many factors could underlie the observed patterns of exonic variation across the 20 regions in our study. In previous work, we evaluated the roles played by founder effects, genetic drift and relaxation of purifying selection in shaping the isolated Shetland genome^8^. While founder effects appear to play a role in the more isolated populations in our study (Shetland, Orkney, full AJ), given the small amount of shared ultra-rare exonic variants per individual in other groups (Table S1) it is unlikely that this is a major force driving the observed regional differentiation for the remaining regions.

To investigate the potential role of genetic drift, we focused on variants found in each of the 20 regions which are also present in the gnomAD database^29^. More specifically, we focus on gnomAD SNPs present in non-Finnish Europeans (NFE) with frequency of 1% ≤ MAF_NFE_ < 5% (generally regarded as rare SNPs) and we assume that this NFE data represents a relatively cosmopolitan European reference group. Each individual in our study carries on average about 1100 such rare variants (Table S1) and a comparison of the MAF of these rare SNPs across all regions as they have diverged from the MAF in gnomAD NFE individuals should mainly reflect the strength of genetic drift in shaping the regional exomes.

Our results show clear evidence of genetic drift in the rare variants found in regional exomes (Fig 3A). Almost half of all known rare SNPs in full AJ are seen at least two-fold more or less frequently in this group compared with the frequency observed in gnomAD NFE, such that 32.0% and 12.7% of SNPs have drifted to lower (down-drift) or higher (up-drift) frequencies, respectively (Table S3). Substantial proportions of rare SNPs also appear to have been subject to genetic drift in the Shetland (down-drift 15.3%, up-drift 3.6%), Orkney (13.7%, 3.1%) and part AJ (10.9%, 4.8%) groups; with more modest effects seen in the Irish (7.9%, 1.2%), Wales N (6.2%, 0.8%) and Wales S (6.1%, 0.5%) groups. The remaining regions show roughly similar proportions of drifted rare SNPs (4.9% on average), with some degree of genetic drift evident even in the relatively cosmopolitan London region (4.2% and 0.2%, respectively), which could be regarded as a baseline level of drift in UK populations compared to the gnomAD Non-Finnish European combined reference population.

**Figure 3.**
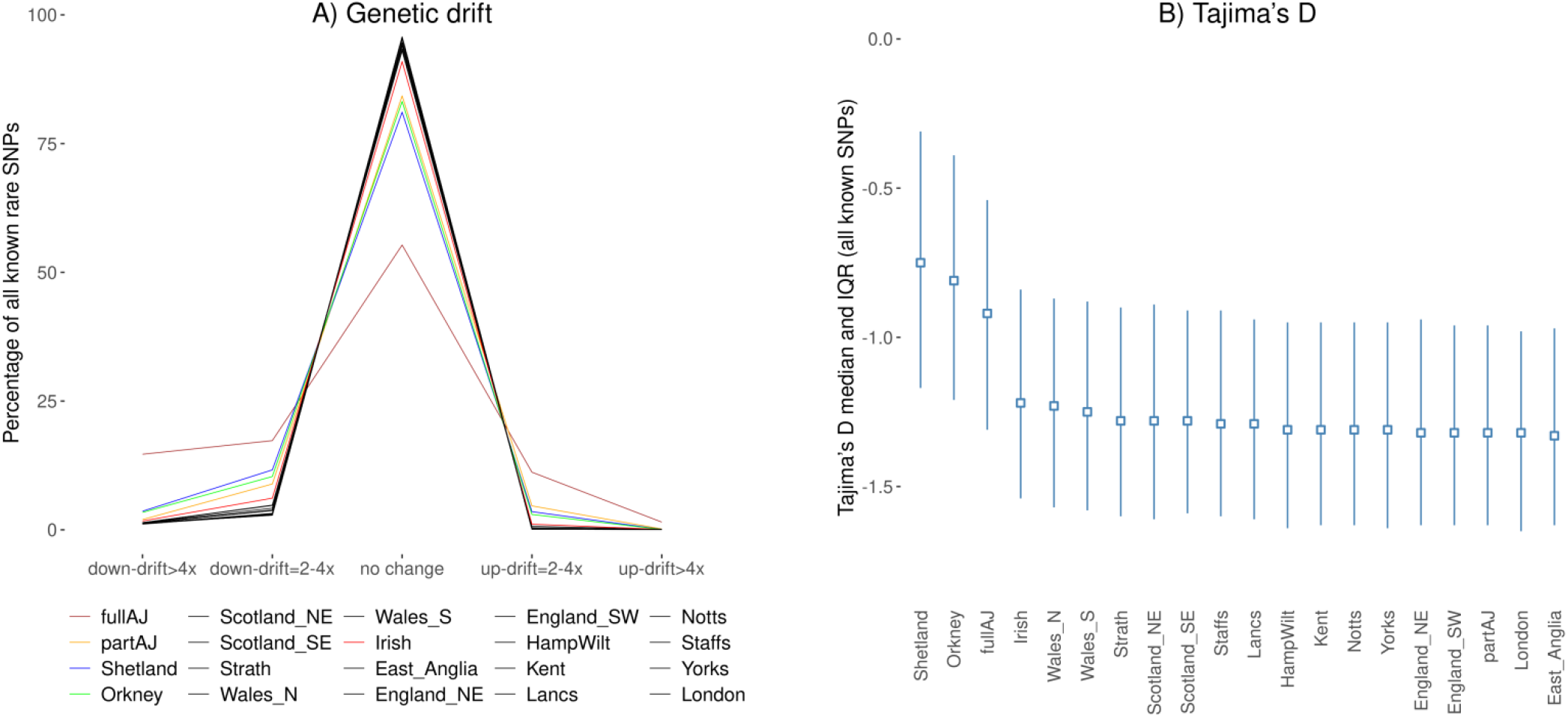
Genetic drift and Tajima’s D within regional populations of the UK and Ireland. **A)** The rare known SNP variants in each of the 20 groups are split into five categories based on the fold change in their regional frequency compared to the frequency with which they are observed in gnomAD NFE individuals as shown on the x-axis. The y-axis represents variants in each of the five categories as a proportion of all known rare SNPs (per group); **B)** Tajima’s D median IQR values in the 20 regions (sorted by median) based upon all known SNP variants found within each regional group.

The roles played by the degree of geographical and/or cultural isolation in shaping the regional exomes was further evidenced by the results of Tajima’s D analysis^30^ (Methods) for all known SNPs present across the 20 regions (Fig 3B, Table S4). Tajima’s D values close to zero are considered as evidence for the neutrality, while negative values may reflect a high number of rare alleles due to population growth and/or purifying selection and positive Tajima’s D value indicate a high number of alleles shared within the population^31^, which can be indicative of relaxation of purifying selection due to relative isolation and reduced effective population size.

The two Northern Isles populations, Shetland and Orkney, exhibit the highest Tajima’s D values, corresponding to their geographically isolated nature and the relatively constant population size^32^ in the recent past. The other isolated population, the full AJ, although going through a tight population bottleneck in the relatively distant past, has experienced a steady population growth since then leading to lower Tajima’s D values compared to the Northern Isles. Part AJ, due to their recent admixture with the general UK population, which in the context of Tajima’s D analysis can be considered as explosive population growth, are virtually indistinguishable from the mainland UK populations. The ranking of the remaining regions based on their Tajima’s D values (Irish, Wales, Scotland and England) corresponds well with the results of our genetic drift analysis (Fig S7).

### Identification of regionally enriched deleterious variants

Based on the observed regional stratification in UKB and the regionally variable influence of genetic drift, we sought evidence for the presence of potentially deleterious exonic variants enriched in particular geographical regions. We conservatively restricted our analysis to variants predicted to affect the coding potential of canonical transcripts, causing stop codon gain, start codon loss, splice donor/acceptor site loss, and frameshifts, as well as missense and splice region variants confidently predicted to be deleterious (CADD score ≥ 30). From the variants identified in these classes we then defined as enriched those found at a regional frequency at least 5 times higher than the frequency observed in gnomAD NFE and attaining statistical significance (Methods). Overall, we discovered at least one enriched and potentially deleterious variant in 14 of the considered 20 UKB regions, summing up to 67 unique variants. These variants are: *i*) enriched in one or more of the UKB regions compared to NFE in gnomAD, *ii*) predicted to be functional, *iii*) implicated in a Mendelian disorder and *iv*) reported in ClinVar^33^ to be pathogenic/likely pathogenic (Methods). The vast majority (95%) are previously known, but extremely rare variants, with 90% of these having gnomAD MAF_NFE_ < 0.0004 (Fig 4).

**Figure 4.**
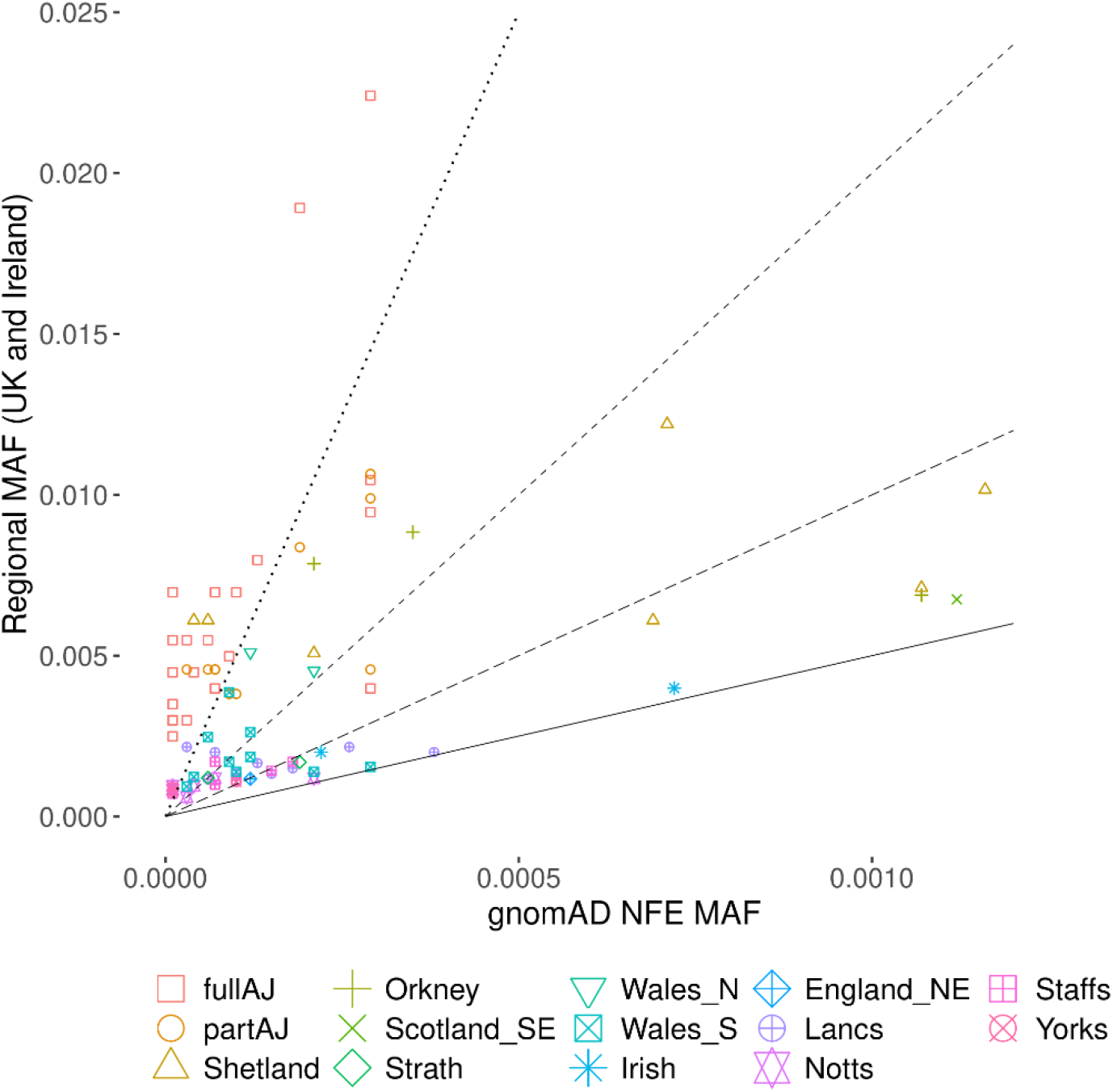
Regionally enriched deleterious variants discovered in the UKB regions of the UK and Ireland. Each of the 67 discovered variants is represented as a point with the frequency at which it is found in gnomAD NFE individuals (x-axis) and its regional frequency (y-axis). Note that, for visual clarity, the two axes are on different scales. To facilitate variant enrichment interpretation, added are four guide lines representing variant regional MAF enrichment of 5 times (solid line), 10 times, 20 times and 50 times (dotted line) compared to gnomAD NFE. Precise enrichment information per each variant is available in the subsequent tables.

### Analysis of the reference AJ group within UKB is instructive

Our analysis revealed 24 enriched and potentially deleterious exonic variants in UKB participants with AJ descent, with 10 of these variants being shared by full and part AJ, while the remaining 14 are seen exclusively in full AJ (Table 1). All of the identified variants exhibit regional MAF (MAF_REG_) at least 10 times higher compared to NFE in gnomAD and more than half have frequencies more than 100 times higher. Most of the identified variants are correlated with health conditions previously reported to be significantly enriched in individuals with Jewish origins^34^ – nine are predominantly AJ diseases, three are mostly found in Sephardi-Mizrahi Jewish and two are common in all Jewish groups (Table 1). In addition, there is a higher incidence of various types of Retinitis pigmentosa among individuals with Jewish heritage^34^, as well as increased risk of developing breast and ovarian cancer among AJ women^35^. The rediscovery of these variants in our UKB analysis supports the effectiveness and accuracy of our approach for identifying deleterious variants enriched in UKB regions.

**Table 1.** Enriched and potentially deleterious variants in Ashkenazi Jewish samples. Conditions found primarily in Ashkenazi Jewish (*), in Sephardi-Mizrahi Jewish (^) and in all Jewish groups (^+^)

| ClinVar Variant<br>Allele ID | Gene | Condition | Region | MAF<sub>REG</sub> | Enrichment<br>(vs MAF<sub>NFE</sub>) |
| --- | --- | --- | --- | --- | --- |
| 98339 | *CPT2* | Carnitine palmitoyltransferase II deficiency* | full AJ | 0.0070 | 474x |
| 20541 | *ADAMTS2* | Ehlers-Danlos syndrome dermatosparaxis type* | full AJ | 0.0055 | 373x |
| 29282 | *MTTP* | Abetalipoproteinaemia* | full AJ | 0.0045 | 305x |
| 361210 | *TECPR2* | Spastic paraplegia 49, autosomal recessive^ | full AJ | 0.0035 | 237x |
| 19617 | *BBS2* | Retinitis pigmentosa | full AJ | 0.0035 | 237x |
| 340082 | *BLM* | Bloom syndrome* | full AJ | 0.0030 | 203x |
| 230781 | *DNAI2* | Ciliary dyskinesia, primary, 9* | full AJ | 0.0030 | 203x |
| 447466 | *ATP13A2* | Kufor-Rakeb syndrome | full AJ | 0.0030 | 203x |
| 244115 | *CHRNE* | Myasthenic syndrome, congenital^ | full AJ | 0.0030 | 203x |
| 18242 | *FKTN* | Walker-Warburg syndrome* | full AJ<br>part AJ | 0.0055<br>0.0046 | 186x<br>155x |
| 15076 | *FAM161A* | Retinitis pigmentosa | full AJ | 0.0025 | 169x |
| 32701 | *BRCA1* | Breast-ovarian cancer, familial 1 | full AJ | 0.0030 | 102x |
| 28743 | *PEX2* | Zellweger syndrome* | full AJ | 0.0045 | 102x |
| 32049 | *GBJ2* | Nonsyndromic hearing loss and deafness | full AJ<br>part AJ | 0.0189<br>0.0084 | 99x<br>44x |
| 286483 | *SLC3A1* | Cystinuria | full AJ<br>part AJ | 0.0070<br>0.0046 | 95x<br>62x |
| 274143 | *EYS* | Retinitis pigmentosa | full AJ<br>part AJ | 0.0055<br>0.0046 | 93x<br>78x |
| 26930 | *F11* | Hereditary factor XI deficiency disease* | full AJ<br>part AJ | 0.0224<br>0.0099 | 76x<br>34x |
| 94265 | *CCDC65* | Primary ciliary dyskinesia | full AJ<br>part AJ | 0.0070<br>0.0038 | 68x<br>37x |
| 19972 | *PCDH15* | Usher syndrome type 1* | part AJ<br>full AJ | 0.0046<br>0.0040 | 62x<br>54x |
| 22168 | *CFTR* | Cystic fibrosis+ | full AJ | 0.0080 | 60x |
| 24364 | *BRCA2* | Breast-ovarian cancer, familial 2 | full AJ<br>part AJ | 0.0050<br>0.0038 | 56x<br>43x |
| 18928 | *HEXA* | Tay-Sachs disease+ | part AJ<br>full AJ | 0.0107<br>0.0105 | 36x<br>36x |
| 415113 | *NCF* | Chronic granulomatous disease^ | full AJ<br>part AJ | 0.0095<br>0.0046 | 32x<br>16x |
| 20180 | *USH1C* | Usher syndrome | full AJ | 0.0040 | 14x |

### Enriched and deleterious rare exonic variants in Scotland

We discovered nine enriched and potentially deleterious variants in the regions of Scotland considered in our analysis. Four of these variants are specific to the Shetland Islands, with one specific variant found in each of the Orkney, Strathclyde and South East Scotland regions. Two variants were also found to be shared across regions of Scotland: a variant associated with Usher syndrome found to be enriched in Shetland and Strathclyde and another associated with Bardet-Biedl syndrome appearing as enriched in both of the Northern Isles populations (Table 2). For each of the identified variants we also computed a range for the predicted regional number of individuals homozygous for the variant (HOM_ALT_ range, Table 2), with the lower bound based on the assumption of random mating of region’s individuals with the whole of the UK and Ireland (with MAF_AVE_ representing the average variant MAF across the 20 regions in our study) and the upper bound based assuming random mating within the region only.

**Table 2.** Enriched and potentially deleterious variants in samples from Scotland.

| ClinVar Variant Allele ID | Gene | Condition | Region | MAF<sub>REG</sub> | Enrichment (vs MAF<sub>NFE</sub>) | MAF<sub>AVE</sub> | HOM<sub>ALT</sub> range |
| --- | --- | --- | --- | --- | --- | --- | --- |
| 33889 | *CLCN1* | Congenital myotonia | Shetland | 0.0061 | 138x | 0.00008 | [0,1] |
| 21837 | *ADGRV1* | Usher syndrome | Shetland | 0.0061 | 104x | 0.00020 | [0,1] |
|  |  |  | Strathclyde | 0.0012 | 21x |  | [1,3] |
| 23042 | *RDH5* | Fundus albipunctatus | Orkney | 0.0088 | 25x | 0.00035 | [0,2] |
| 23943 | *PPT1* | Neuronal ceroid lipofuscinosis 1 | Shetland | 0.0122 | 17x | 0.00067 | [0,3] |
| 21382 | *FANCF* | Fanconi anaemia | Strathclyde | 0.0017 | 8.8x | 0.00026 | [1,7] |
| 20604 | *AIPL1* | Leber congenital amaurosis | Shetland | 0.0061 | 8.8x | 0.00038 | [0,1] |
| 20006 | *ABCG8* | Sitosterolaemia 1 | Shetland | 0.0102 | 8.8x | 0.00124 | [0,2] |
| 16367 | *BBS10* | Bardet-Biedl syndrome | Shetland | 0.0071 | 6.6x | 0.00128 | [0,1] |
|  |  |  | Orkney | 0.0069 | 6.4x |  | [0,1] |
| 176561 | *LOXHD1* | Nonsyndromic hearing loss and deafness | Scotland SE | 0.0068 | 6.0x | 0.00133 | [13,65] |

### Enriched and deleterious rare exonic variants in Wales

We identified nine enriched and potentially deleterious variants in the Welsh groups in UKB, eight of which were specific to South Wales and one shared with individuals born in North Wales (Table 3). The lack of north Wales specific variants is likely to be explained by the almost four-fold smaller sample size for unrelated North Welsh (n=883) individuals in our study compared to their Southern counterparts (n=3239). Furthermore, it is possible that not all eight South Wales variants are truly specific to this region; some may be shared with the neighbouring English regions (e.g. Gloucestershire, Herefordshire, Shropshire and Cheshire), which were not included in our study due to the insufficient number of unrelated UKB individuals in these regions with WES data available.

**Table 3.**
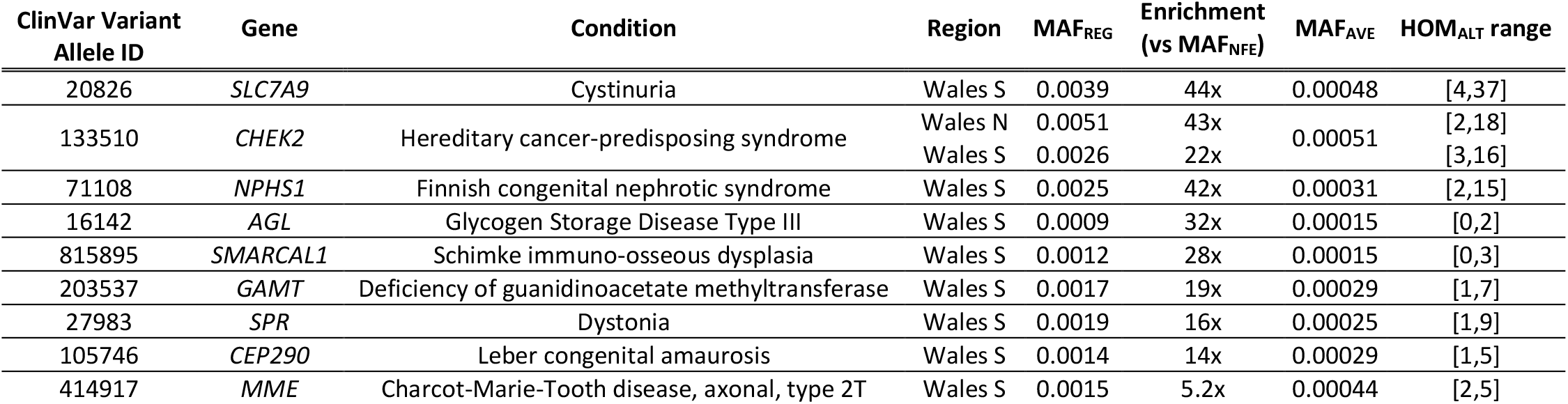
Enriched and potentially deleterious variants in samples from Wales.

### Enriched and deleterious rare exonic variants in England

Our analysis of the WES data from individuals born in the ten English regions discovered 22 enriched and potentially deleterious variants (Table 4). Apart from a single variant found to be enriched in the North East England region (in the *PNP* gene), all of the remaining 21 variants were identified in four neighbouring regions: Lancashire, Staffordshire, Nottinghamshire and Yorkshire. In addition to variants specific to each of these regions, we also identified three variants (in the *COL7A1, F11* and *COL4A4* genes) as shared between two of these regions and one variant (*ALMS1* gene) shared by individuals born in Lancashire, Staffordshire and Nottinghamshire.

**Table 4.**
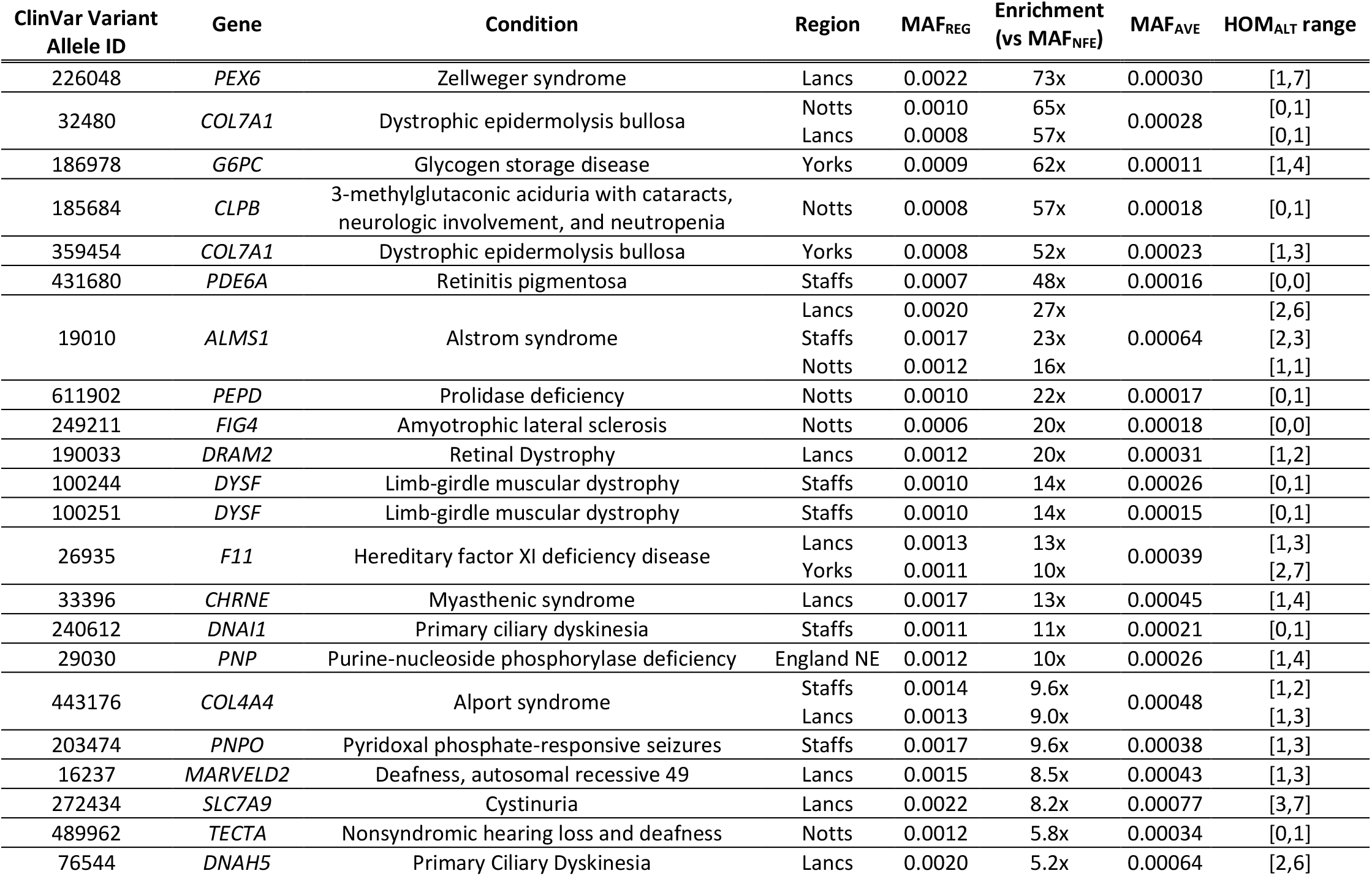
Enriched and potentially deleterious variants in samples from England.

### Enriched and deleterious rare exonic variants in Ireland

The analysis of the 2005 unrelated UKB individuals who self-identify as Irish and were born in either Northern Ireland or the Republic of Ireland resulted in identification of two enriched and potentially deleterious variants (Table 5).

**Table 5.** Enriched and potentially deleterious variants in self-identified Irish individuals born on the island of Ireland.

| ClinVar Variant Allele ID | Gene | Condition | Region | MAF<sub>REG</sub> | Enrichment (vs MAF<sub>NFE</sub>) | MAF<sub>AVE</sub> | HOM<sub>ALT</sub> range |
| --- | --- | --- | --- | --- | --- | --- | --- |
| 401365 | SPG7 | Hereditary spastic paraplegia | Ireland | 0.0020 | 9.0x | 0.00027 | [4,28] |
| 31343 | FMO3 | Trimethylaminuria | Ireland | 0.0040 | 5.5x | 0.00095 | [26,110] |

One reason for the relatively smaller number of enriched variants found in Ireland compared to other mainland UKB regions may be the different sample selection criteria -in contrast to our requirement for individuals in England, Scotland and Wales to exhibit very similar genetic ancestry based on a principal components analysis of the UKB whole-genome SNP array genotypes, the Irish participants were selected only based on self-identification as Irish and being born in Northern Ireland or the Republic of Ireland (Methods). As a result, it is possible that our sample of Irish individuals contains some with non-Irish ancestry, e.g. in the process of selecting Irish individuals we have identified and excluded six participants with AJ heritage. Another factor might be the relatively low number of Irish participants with available WES data. The analysed 2005 unrelated individuals represent the whole population of Ireland (about 7 million), thus inhibiting identification of potential within-Ireland differentiating signal(s).

### Biomedical implications of regionally enriched deleterious variation

The relatively high genomic homogeneity in individuals with Ashkenazi Jewish heritage has been well established and targeted genetic screens for variants implicated in various Mendelian disorders have been adopted world-wide. Enabled by the large-scale WES data made available by UK Biobank, we have demonstrated the existence of analogous deleterious variant enrichment within various geographical regions of the UK and Ireland (Tables 2-5).

While these data highlight single variants with large impact (e.g., the *FMO3* and *SPG7* gene variants in Irish individuals, the *SLC7A9* variant in South Wales, the *LOXHD1* variant in South East Scotland and the *CHEK2* variant in Welsh individuals), they also provide disorder-centric view (e.g., Cystinuria with between 7 and 44 individuals predicted to be homozygous for the enriched causative variants, Primary ciliary dyskinesia with two to seven individuals, Glycogen storage disease with one to six individuals, Leber congenital amaurosis with one to six individuals, Dystrophic epidermolysis bullosa with one to five individuals), as well as regional aggregates (e.g., between 30 and 138 Irish individuals predicted to be homozygous for the regionally enriched deleterious variants, 14 to 99 South Wales homozygous individuals, 13 to 65 South East Scotland individuals, 13 to 42 Lancashire individuals, etc.). It should be noted that the reported numbers of predicted individuals homozygous for the regionally enriched deleterious variants are a clear underestimate of the number of individuals potentially affected by the corresponding condition due to the genetic and locus heterogeneity of the disorders, which is not taken into consideration in our calculations.

A deleterious variant causing a frameshift in the *OBSL1* gene (chr2:219568063:G>GT, c.1273dup, p.T425fs, rs762334954) was found to be regionally enriched in the Northern Isles of Scotland (Orkney and Shetland) and the geographically distant Wales. However, upon closer examination the variant also appears to be measurably enriched in other UKB regions as well, but failing to meet our stringent enrichment criteria there (Table 6). This variant has been previously reported to be associated with the 3-M syndrome^36^, an extremely rare autosomal recessive primordial growth disorder, characterised by distinct facial features, radiological abnormalities, normal intelligence and final adult height in the range of 115 to 150 cm. The exact prevalence of this disorder remains unclear, with around 200 reported cases world-wide as of 2012 since the first published report in 1975, but predicted to have increased substantially with the greater awareness of the disorder and increased availability of genetic testing^37^. To estimate the practical impact of the elevated frequency of the *OBSL1* variant, we considered its effect in each UKB region separately. The variant exhibits regional genetic prevalence of individuals homozygous for it (computed as MAF_REG_^2^) of 1/16k (∼1500 times higher than gnomaAD NFE individuals) in Orkney, 1/39k (∼600 times higher) in Shetland, 1/49k (∼500 times higher) in North Wales and 1/518k (∼45 times higher) in South Wales. Assuming random mating within regions, it is expected there will be 1.4, 0.6, 1.4 and 5.2 homozygous individuals affected by the condition in the Orkney, Shetland, North Wales and South Wales regions, respectively. Given the average MAF=0.000467 in the remaining UKB regions, a genetic prevalence of 1/4.6m (∼5 times higher than NFE) can be expected assuming random mating, which translates to 10.7 individuals affected by the condition across these regions. Overall, due to the regionally elevated frequency of the *OBSL1* variant, we estimate that 19 individuals across the UK and Ireland could be affected by the 3-M syndrome due to being homozygous for this variant.

**Table 6.**
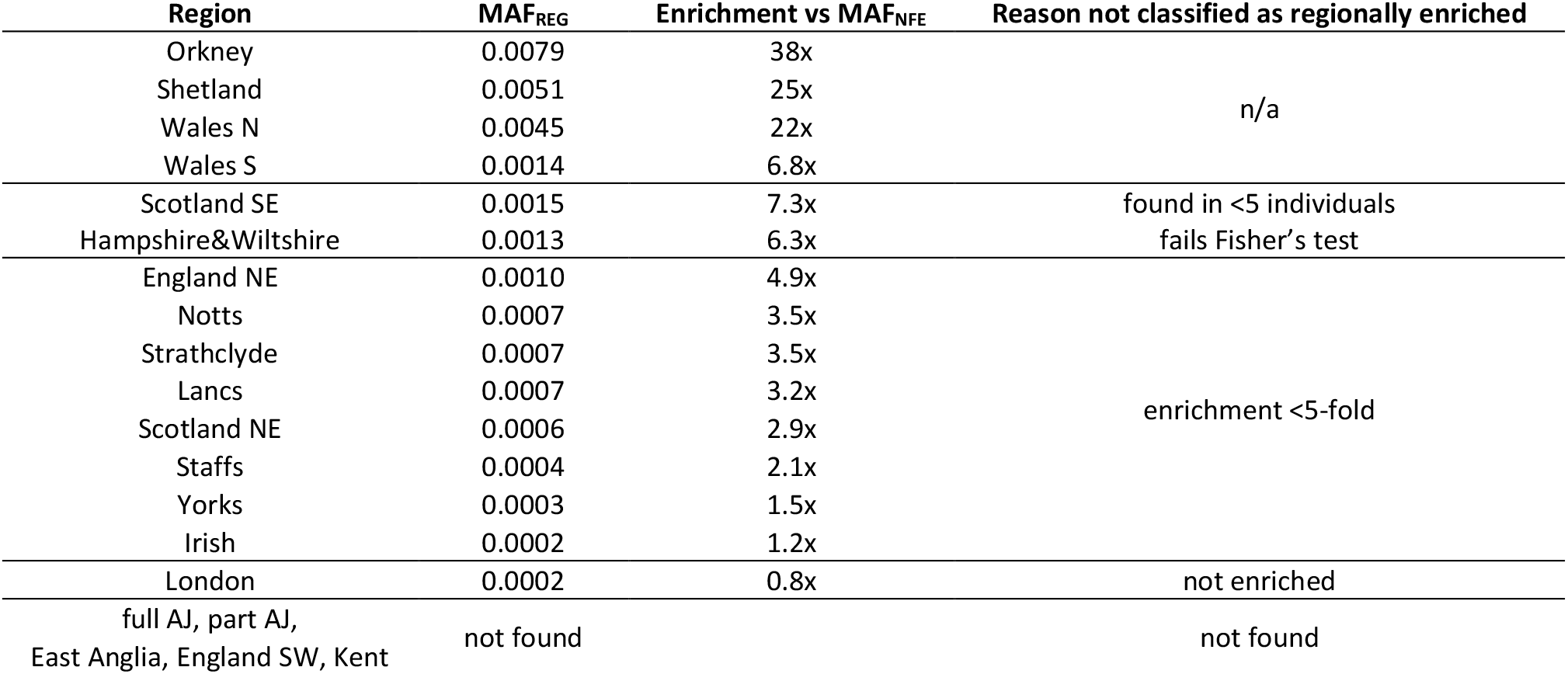
The enriched and potentially deleterious frameshift variant in the *OBSL1* gene (ClinVar Variant Allele ID: 192467, Three M syndrome 2).

## Discussion

The work described here is based on the UKB WES data for the first ∼200,000 participants released in October, 2020^24^. We complemented these data with our own WES data for more than 4,000 individuals from Shetland and Orkney (the Northern Isles) in Scotland, which were generated by the same provider (Regeneron Genetics Center) using the same Original Quality scores Functional Equivalence (OQFE) protocol as for the UKB dataset, to compile a combined dataset of 44,696 unrelated individuals born in various geographical regions (Methods). Based on the availability of sufficient numbers of UKB individuals with WES data, we divided individuals into 18 geographical regions: two representing the Northern Isles, three Scottish regions, two Welsh regions, ten English and one Irish region, consisting of individuals who self-identify as Irish and were born in either Northern Ireland or the Republic of Ireland (Methods). Additionally, based on the available genetic and genomic data, we identified the presence of individuals with Jewish heritage in the UKB dataset. We present several lines of evidence supporting our hypothesis these are mainly individuals of Ashkenazi Jewish (AJ) origin and which we define as full and part AJ individuals based on the degree of their Ashkenazi Jewish heritage, for a total of 20 genetic groups considered in our study.

Previous research showed a remarkable correlation between genomic variation and the geography of the UK and Ireland^5,6,26^. Here, we demonstrate for the first time that this signal is largely preserved in protein-coding exons in spite of these regions general intolerance to variation. Our results show a clear separation of the isolated Shetland and Orkney Isles populations as well as AJ individuals from each other and the other regions, and we recapture patterns consistent with reports of a Welsh-English-Scottish-Irish cline from previous studies based on genome-wide genotyping arrays^5,6^. Our results also revealed a distinction between individuals born in North and South Wales, and some level of distinctiveness exhibited by South East Scotland individuals. These findings should be considered when performing association studies, where accounting for population structure is of crucial importance. The geographical variation in genotypes in the UKB has been documented before in the context of GWAS studies^38,39^, our results suggest similar care must be taken in future exome-wide association studies based on UKB data.

Our analyses suggest that the observed regional structure is driven mainly by two factors. Firstly, based on the frequency with which variants are found in our data compared to their frequency in Non-Finnish Europeans (NFE) in gnomAD, we found evidence of genetic drift present in all 20 UKB regions. The proportion of SNP loci exhibiting substantial genetic drift in each group correlates well with the assumed degree of geographical/cultural/social isolation, being highest in AJ and the Northern Isles, followed by Irish, Welsh, Scottish and English regions, but noticeable even in the cosmopolitan London region (Fig 3A, Table S3). An IBD-based metric of isolation in mainland UK, termed “preference of local connectivity”, has been described previously^26^ and found to be inversely correlated with population density. The results reported by Naseri et al.^26^ are concordant with our genetic drift findings, showing people from Greater London with weakest local connectivity, followed by moderate connectivity in central England and strong local connectivity, even in major cities, in Scotland and Wales. Secondly, a Tajima’s D analysis based on all known autosomal SNPs revealed the roles played by population growth and purifying selection in shaping the British and Irish exomes. Due to their relatively constant population size and geographically isolated nature, the two Northern Isles populations (Shetland and Orkney) exhibit the highest Tajima’s D values (Fig 3B) indicative of the largest relaxation of purifying selection among the 20 regions. Despite going through a severe genetic bottleneck in the past, the AJ group has experienced steady population growth since, resulting in weaker relaxation of purifying selection in our full AJ group compared to the Northern Isles and virtually no difference between the admixed part AJ and the remainder of the UK population. The ranking of the remaining regions based on their Tajima’s D values (Irish, Wales, Scotland and England) corresponds well with the results of our genetic drift analysis and the assumed degree of regional isolation.

The most important practical implication of the observed regional variation, driven by genetic drift and relaxation of purifying selection as a result of various geographical/cultural/social barriers, is the elevated frequency of some otherwise rare variants with proven links to human health. In our study, we focused on significantly enriched exonic variants otherwise rarely or not found in a healthy control dataset (gnomAD MAF_NFE_ < 1%), predicted to be functional and previously reported as implicated in Mendelian disorders (Methods). Applying stringent filtering criteria, we found 67 such unique variants enriched in one or more of the considered groups compared to NFE in gnomAD with 95% of these variants being present, but extremely rare in gnomAD (90% of them with gnomAD MAF_NFE_ < 0.0004, i.e. 1/2500, Fig 4) and the remaining 5% not found in any gnomAD population. Our analyses based on random mating suggest that the regionally enriched and potentially deleterious variants can be expected to result in a significant number - tens or hundreds - of individuals affected by a recessive genetic medical condition, which highlights the importance of future research into regional variation across the UK and Ireland, to inform effective genetic screening and counselling.

There are several key issues that need to be addressed in order to provide a more robust regional variation knowledge base. Firstly, our results are based on the analysis of 44,696 unrelated UKB individuals with WES data available, which constitutes less than 0.1% of the overall UK and Republic of Ireland population. In order to accurately estimate the regional frequency of these extremely rare deleterious exonic variants, a deeper, wider, more ethnically diverse and as random as possible sampling would be beneficial. This point is further illustrated by the observation that for five of the six regions in our study with no enriched variant discovered - East Anglia, Kent, England SW, Scotland and Hampshire&Wiltshire - the number of unrelated individuals included in our study is below the median number of participants for the remaining 14 regions. It can be expected that the release of the next UKB WES tranche of 500k individuals would alleviate this problem to a certain extent, as well as it increasing the breadth of regional coverage. The availability of such information will help in better understanding of the geography of the variant enrichments. For example, are the variants discovered to be enriched in South Wales region-specific, or are some of them also enriched in the neighbouring Welsh Marches (e.g. Gloucestershire, Herefordshire, Shropshire and Cheshire, in England), for which at the moment we lack sufficient UKB participants with WES data available? All these facts highlight an important limitation of our study as is, namely no firm conclusions can be drawn by comparing the numbers of discovered enriched variants across regions (as coverage is so variable). In addition, despite the inclusive efforts embedded in the UKB recruitment, there is participation bias towards older (median age at assessment = 58), healthier individuals from more economically affluent areas (median Townsend deprivation index score = -2.2) who self-identify as “White” (93.7%)^24,40^, and who live near the 22 UKB assessment centres, 17 of which were in England and none of which were in Northern Ireland, making the dataset not fully representative of the overall UK population.

Secondly, our strategy for identifying regionally enriched deleterious variants favours specificity over sensitivity. For example, our decision to consider only missense variants with CADD score ≥ 30 leads to omitting the rs76763715 variant, a common pathogenic variant reported in the homozygous and compound heterozygous state in individuals with Gaucher disease (type I). This condition exhibits a higher prevalence among individuals with Ashkenazi Jewish heritage and in our data the variant is 13 times more frequent in AJ individuals compared to London, but was excluded due to a CADD score of 24. Next, our chosen threshold of considering as enriched only regional variants observed with frequency at least 5 times higher compared to Non-Finnish Europeans is not necessarily optimal (e.g. the 3-M syndrome variant); a lower, or even no such threshold, may be more relevant from a medical perspective, highlighting the crucial importance of close collaboration between genetic scientists, clinicians, stakeholders and policy-makers. Further, in assessing the predicted effect of the discovered enriched variants, we retained only variants with substantial evidence of being pathogenic/likely pathogenic available as reported by ClinVar; however only about 20% of all enriched variants identified in our study were found with any pathogenicity annotation in ClinVar. Finally, in estimating the practical impact of the regionally enriched variants we focused on calculations of the genetic prevalence of individuals homozygous for these variants. This leads to a clear underestimation of individuals affected by the corresponding medical condition, due to the genetic and locus heterogeneity of the Mendelian recessive disorders and highlights the need of an additional, disorder-centric, systematic investigation of the regional frequency of all other known related variants. As a result of all these factors, we view the reported 67 regionally enriched and potentially deleterious variants and their practical implications just as an illustration of the potentially much larger set of medically relevant and clinically important regional variants in the UK and Ireland.

## Methods

### Ethics statement

All participants in the Viking Health Study—Shetland (VIKING) gave written informed consent for broad ranging health and ancestry research including, whole genome/exome sequencing and the study was given a favourable opinion by the South East Scotland Research Ethics Committee (REC Ref 12/SS/0151). All participants in the Orkney Complex Disease Study (ORCADES) gave written informed consent for broad ranging health and population research, including sequencing and the study was approved by Research Ethics Committees in Orkney, Aberdeen (North of Scotland REC), and South East Scotland REC, NHS Lothian (reference: 12/SS/0151).

### Participant selection

The Viking Health Study - Shetland (VIKING)^9^ and Orkney Complex Disease Study (ORCADES)^25^ are family-based, cross-sectional studies that seek to identify genetic factors influencing cardiovascular and other disease risk in the population isolates of the Shetland and Orkney Isles in northern Scotland. These studies are now subsumed, along with VIKING II, under the Viking Genes umbrella (https://www.ed.ac.uk/viking). 2105 participants were recruited to VIKING between 2013 and 2015, most having at least three grandparents from Shetland, while 2078 participants were recruited to ORCADES between 2005-2011, most having three or four grandparents from Orkney, the remainder with two Orcadian grandparents. Fasting blood samples were collected and many health-related phenotypes and environmental exposures were measured in each individual.

### Northern Isles

As part of Viking Genes (https://www.ed.ac.uk/viking), more than 4000 individuals from the Shetland and Orkney islands (the Northern Isles of Scotland) were selected for whole exome sequencing. Since the focus of this work is on regional variation, we utilized the rich genealogical information collected as part of Viking Genes and from the 2134 Shetland participants with WES data we selected for further analysis the 1454 individuals with all four grandparents also born on the Shetland archipelago. To alleviate familial effects and to obtain a more representative snapshot of the regional variation, we identified related individuals up to first cousins once removed and closer and equivalents (PLINK v1.90b4^41^; pi_hat >= 0.0625) and generated the maximum unrelated set (using PRIMUS v1.9.0^42^) of 492 unrelated Shetlanders with all four grandparents born on the Shetland archipelago. Similarly, from the 2092 Orkney participants with WES data we selected a maximum unrelated set of 509 unrelated Orcadians with all four grandparents born on the Orkney archipelago.

### UKB

The UK Biobank Exome Sequencing Consortium (UKB-ESC) is a private-public partnership between the UKB and eight biopharmaceutical companies that will complete the sequencing of exomes for all ∼500,000 UKB participants^24^. To explore the regional variation, we considered geographical region of birth based on historic counties and selected only those regions for which there were at least 500 UKB participants with WES data available, using the 200k WES UKB tranche (released in October 2020) after excluding all participants who withdrew their consent. Since grandparents’ birthplace information is not available for the UKB participants, in order to focus on the regional variation in the UKB data we selected only participants who self-identify as “White British” (UKB field: 21000), exhibit very similar genetic ancestry based on a principal components analysis of the UKB whole-genome SNP array genotypes (UKB field: 22006) and who were born outside large metropolitan areas in the corresponding region. The participants satisfying the above criteria for each region were then evaluated for relatedness and the maximum unrelated set per region generated as for the Northern Isles cohorts.

The three **Scotland** regions we included in our study are: **Scotland North East** (Aberdeen, Aberdeenshire, Kincardineshire, Angus, Banffshire, Dundee, Fife, Perthshire, Kinross-shire, Clackmannanshire, Stirlingshire, Moray) with 1680 unrelated individuals, **Scotland South East** (East Lothian, Midlothian, West Lothian, Selkirkshire, Berwickshire, Roxburghshire, Peeblesshire; excluding Edinburgh) with 667 unrelated individuals and the south-western region of **Strathclyde** (Lanarkshire, Renfrewshire, Dunbartonshire, Ayrshire; excluding Glasgow) with 2077 unrelated individuals (Table S1).

The two **Wales** regions we included in our study are: **Wales North** (Anglesey, Caernarfonshire, Merionethshire, Montgomeryshire, Flintshire, Denbighshire) with 883 unrelated individuals and **Wales South** (the remaining part of Wales; excluding individuals born in a 5 mile radius area centred on Cardiff) with 3239 unrelated individuals.

The ten **English** regions we included in our study are: **England North East** (Northumberland and Durham; excluding individuals born in a 15 mile radius area centred on Newcastle-upon-Tyne) with 2982 unrelated individuals, **Yorkshire** (excluding individuals born in a 5 mile radius areas centred on Kingston-upon-Hull and Doncaster and 15 mile radius areas centred on Leeds, Bradford and Sheffield) with 3276 unrelated individuals, **Lancashire** (excluding individuals born in two 15 mile radius areas centred on each of Liverpool and Manchester) with 3007 unrelated individuals, **Nottinghamshire** with 4192 unrelated individuals, **Staffordshire** (excluding individuals born in a 10 mile radius areas centred on Birmingham and Wolverhampton) with 3526 unrelated individuals, **East Anglia** (Norfolk and Suffolk) with 923 unrelated individuals, **Hampshire and Wiltshire** (excluding individuals born in a 5 mile radius areas centred on Portsmouth and Southampton) with 1925 unrelated individuals, **Kent** (excluding individuals born in a 17 mile radius area centred on the City of London) with 1327 unrelated individuals and **England South West** (Cornwall and Devon) with 1412 unrelated individuals. We also included **Central London** (individuals born in a 10 mile radius area centred on the City of London) with 8913 unrelated individuals, which given the cosmopolitan nature of the capital are expected to serve as a useful control in terms of regional variation.

We have also included in our analysis as **Irish** a group of 2005 unrelated individuals who self-identify as Irish (UKB field: 21000) and born in either Northern Ireland or the Republic of Ireland (UKB field: 1647).

The last group of UKB participants included in our study are individuals of **Ashkenazi Jewish (AJ)** ancestry, split into **full AJ** (1004 unrelated individuals) and **part AJ** (657 unrelated Individuals).

### Sequencing, mapping and variant calling

The WES sequencing, read mapping and variant calling for the ∼200k UKBB participants was performed following the OQFE protocol^24^. The WES sequencing, read mapping and variant calling for the individuals from the Northern Isles (Shetland and Orkney) was performed at the same sequencing facility using the same sequencing and data processing protocols as for the UKBB participants. The starting point for our analyses were the project VCF files generated by the OQFE protocol.

### Variant QC and annotation

Using the unrelated individuals selected for each of the 20 regions described above, we generated 20 regional VCFs by extracting from the corresponding project data only variation present in the particular region, excluding non-variant sites and sites with more than 10% missing genotypes. The regional VCF was decomposed and normalized and the remaining missing genotypes (‘./.’) were set to homozygous reference genotype (‘0/0’). All variants in the 100bp flanking regions outside the capture region were excluded, as well as variants in low-complexity regions based on sdust^43^ or failing the filtering criteria in gnomADg v3.1.1^29^.

Further, any individual SNPs with read depth (DP) < 7, genotype quality (GQ) < 10, heterozygous SNPs with variant allele frequency (VAF) < 0.15 or VAF > 0.85, homozygous SNPs with VAF < 0.85 and any SNPs in windows with problematic gnomADg v3.1.1 coverage (defined as 10bp windows centred on any base with coverage < 10x) were excluded. Similarly, any individual INDELs with DP < 10, GQ < 10, heterozygous INDELs with VAF < 0.2 or VAF > 0.8 and homozygous INDELs with VAF < 0.8, are excluded. The information about the number of variants filtered at each step is reported in Table S5.

The variants in the resulting VCF were annotated with their predicted functional effect using VEP^44^ (v102) including annotation of each variant with its MAF as reported in gnomADg v3.1.1. Lastly, we excluded any variant if it has not been observed in gnomADg v3.1.1 but is detected in the regional VCF with MAF ≥ 10%.

### MDS and UMAP analysis

Our MDS and UMAP analyses are based on a dataset of 10,001 unrelated individuals (492 Shetlandic, 509 Orcadian and 500 randomly chosen individuals from the remaining 18 regions) and on their autosomal coding SNP variation. We considered only SNPs also found in gnomAD in order to alleviate the potential effect of false positive variation in our data. From the assembled VCF, we excluded all variants in regions with known long-range high linkage disequilibrium^45,46^, as well as all singleton variants. From the remaining 512,327 SNPs we selected only biallelic variants with observed AF in our dataset of less than 5%. The selected 465,647 SNPs were screened and pruned based on further linkage disequilibrium evidence (r^2^<0.02; PLINK 1.90b4 with --indep-pairwise 500 5 0.2) resulting in a final set of 401,895 markers which we used as an input for our Multi-Dimensional Scaling (MDS) analysis performed with PLINK (--cluster --mds-plot 50, Fig S3A). The UMAP analysis (e.g. Fig 1A) was performed on the top 20 MDS dimensions using the general_umap_script.py^18,47^.

### Unrooted NJ phylogenetic tree of the 20 regions

A commonly used metric for evaluating the similarity between populations is the Weir and Cockerham *F*_*ST*_ fixation index^48^, which is a measure of population differentiation due to genetic structure and represents the relative amount of genetic variance between populations compared to the total genetic variance within these populations^49^. We calculated the pair-wise *F*_*ST*_ distances among the 20 regions based on non-singleton known SNPs with MAF < 5% in our chosen set of 10,001 unrelated individuals using the--weir-fst-pop option in VCFtools^50^ (v0.1.13) with window size of 1Mb and window step of 250kb. For constructing the phylogenetic tree we used PHYLIP^51^ (v3.697) in Neighbour-Joining (NJ) mode using full AJ as outgroup and randomized the region input order. The computed tree is visualized with iTOL^52^.

### Tajima’s D analysis

Tajima’s D analysis of the 492 Shetland, 509 Orkney and the 500 randomly selected unrelated individuals for the remaining 18 regions was performed using VCFtools (v0.1.13) using the--TajimaD option and sliding windows of size 1Mb. The analysis was based on the regions’ known SNPs (i.e., found with passing quality in the gnomAD dataset) identified in the 22 autosomal chromosomes. For each region, we computed the median Tajima’s D value and it’s IQR by aggregating the results observed for the ∼500 individuals in the ∼3000 genomic windows of size 1Mb, excluding any window with no SNPs present.

### Regionally enriched deleterious variation

In order to increase our discovery power, we based our search for regionally enriched potentially deleterious variants on the full subsets of unrelated individuals per each region (Table S1). To focus on functional variation, our analysis is restricted to variants on canonical transcripts with VEP predicted stop gained, start lost, splice donor/acceptor site (the last 2bp at each end of the intron), stop lost and frameshift effects, as well as missense and splice region variants (1-3 bp into an exon or 3-8 bp into an intron) for which we imposed the additional criterion of having CADD score ≥ 30. From these variants, we selected as enriched variants those that exhibit regional frequency at least 5 times higher than the variant’s frequency observed in the genomes of 34,029 Non-Finnish Europeans (NFE) in gnomAD^29^ (v3.1.1) and the enrichment being statistically significant (Fisher’s Exact Test, Bonferroni adjusted for multiple testing). Variants with regional MAF ≥ 5% (MAF_NFE_ ≥ 1%) are excluded as less likely to be implicated in Mendelian disorders, as well as variants seen in less than 5 unrelated individuals within region, in order to focus on regional, rather than familial variation. Lastly, we screened the regionally enriched variants against ClinVar^33^ and selected only those reported in ClinVar to be pathogenic/likely pathogenic with criteria provided, multiple submitters and no conflicting interpretation (at least 2 star variants) and further manually curated by us to select only variants explicitly reported in peer-reviewed publication(s).

## Acknowledgements

The Viking Health Study – Shetland (VIKING) was supported by the MRC Human Genetics Unit quinquennial programme grant “QTL in Health and Disease”. DNA extractions and genotyping were performed at the Edinburgh Clinical Research Facility, University of Edinburgh. We would like to acknowledge the invaluable contributions of the research nurses in Shetland, the administrative team in Edinburgh and the people of Shetland.

The Orkney Complex Disease Study (ORCADES) was supported by the Chief Scientist Office of the Scottish Government (CZB/4/276, CZB/4/710), a Royal Society URF to J.F.W., the MRC Human Genetics Unit quinquennial programme “QTL in Health and Disease”, Arthritis Research UK and the European Union framework program 6

EUROSPAN project (contract no. LSHG-CT-2006-018947). DNA extractions were performed at the Edinburgh Clinical Research Facility, University of Edinburgh. We would like to acknowledge the invaluable contributions of the research nurses in Orkney, the administrative team in Edinburgh and the people of Orkney.

We acknowledge support from the MRC Human Genetics Unit programme grant, “Quantitative traits in health and disease” (U. MC_UU_00007/10). Finally, we thank UK Biobank, approved under project 19655.

For the purpose of open access, the author has applied a Creative Commons Attribution (CC BY) licence to any Author Accepted Manuscript version arising from this submission.

## Data Availability

For ORCADES and VIKING, there is neither Research Ethics Committee approval, nor consent from individual participants, to permit the open release of the individual level research data underlying this study. The datasets analysed during the current study are therefore not publicly available. Instead, the research data and/or DNA samples are available from on reasonable request, following approval by the QTL Data Access Committee and in line with the consent given by participants. Each approved project is subject to a data or materials transfer agreement (D/MTA) or commercial contract. The UK Biobank genotypic data used in this study were approved under application 19655 and are available to qualified researchers via the UK Biobank data access process.

