## Supplemental Data for "Regionally enriched rare deleterious exonic variants in the UK and Ireland"

### Supplementary Figures

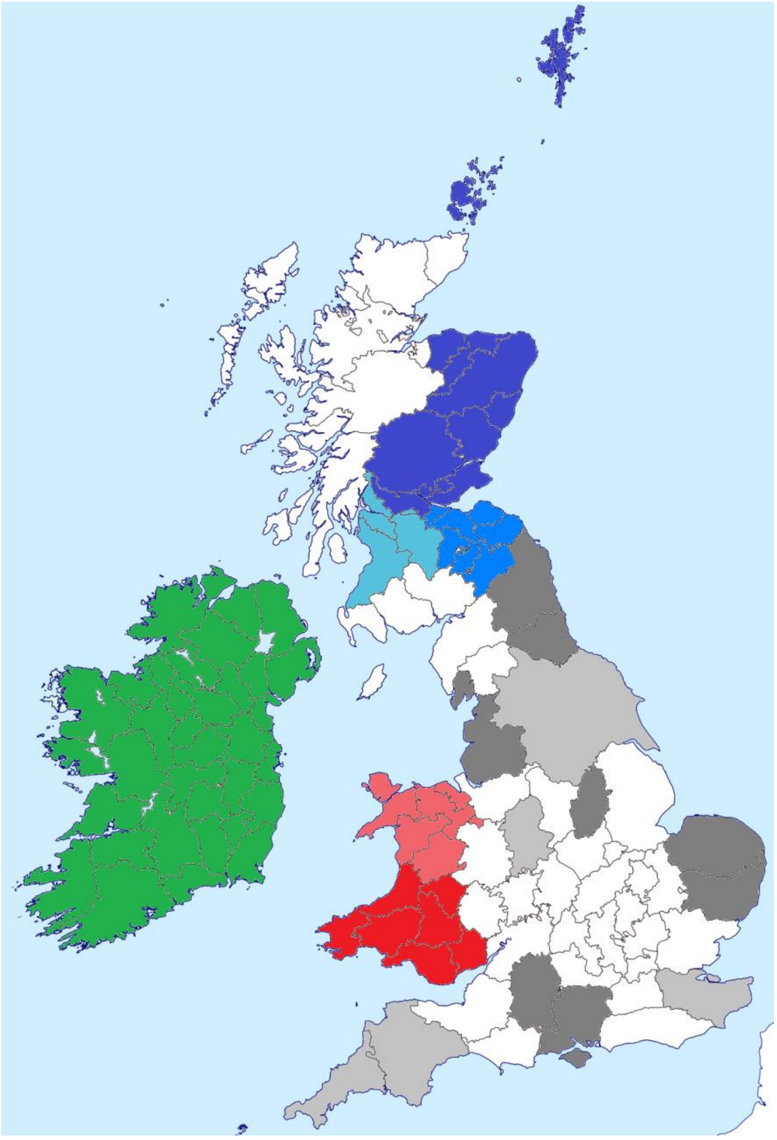

**Figure S1.** Regions of origin of the participants in this study

The Shetland and Orkney (the Northern Isles of Scotland) participants are obtained from the Viking Genes project (https://www.ed.ac.uk/viking). The three mainland UKB Scotland regions (in blue) we included in our study are Scotland North East, Scotland South East and the south-western region of Strathclyde. The two Wales regions (in red) we included in our study are Wales North and Wales South. The ten English regions (in grey) we included in our study are England North East, Yorkshire, Lancashire, Nottinghamshire, Staffordshire, East Anglia, Hampshire and Wiltshire, Kent and England South West. We also included Central London (individuals born in a 10 mile radius area centred on the City of London). Irish participants were selected based in self-identification as Irish and born in either Northern Ireland or the Republic of Ireland (in green). The last group of UKB participants included in our study are individuals of Ashkenazi Jewish (AJ) ancestry, split into full and part AJ, regardless of their geographical region of origin.

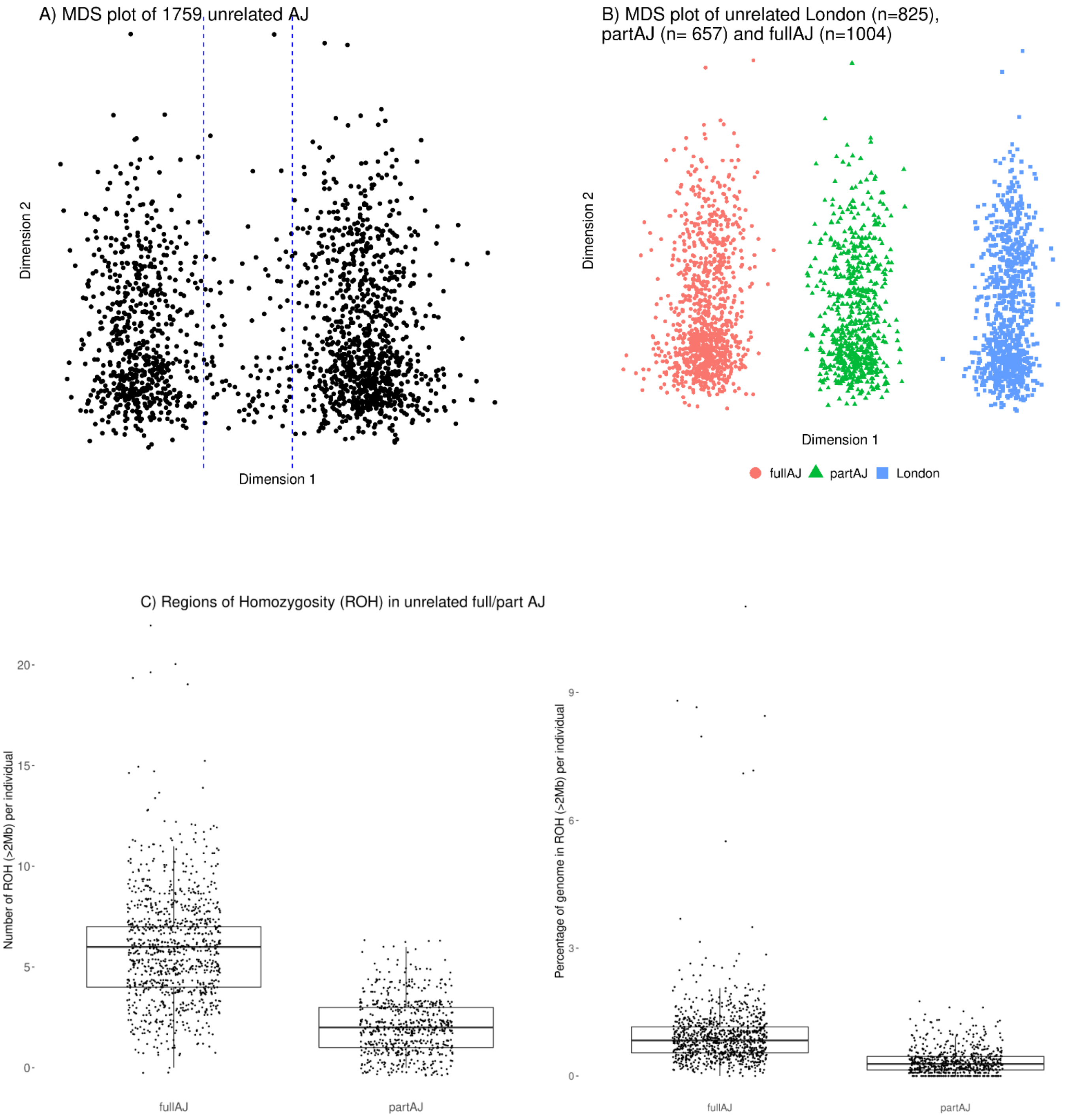


**Figure S2.** Analyses of unrelated Ashkenazi Jewish individuals' heritage

**A)** MDS plot based on biallelic, non-singleton and linkage-disequilibrium (LD) pruned known SNPs (MAF > 1%) in 1759 unrelated AJ individuals reveals two main clusters speculated to represent different level of admixture between AJ and general UK population; **B)** MDS plot of the selected individuals from the two main AJ clusters (left and right from the vertical dashed blue lines in panel A) coupled with 825 unrelated Londoners based on biallelic, non-singleton and LD pruned known SNPs (MAF > 5%) ; **C)** Analyses confirm full AJ individuals exhibit higher number/larger proportion of genome in ROH compared to part AJ.

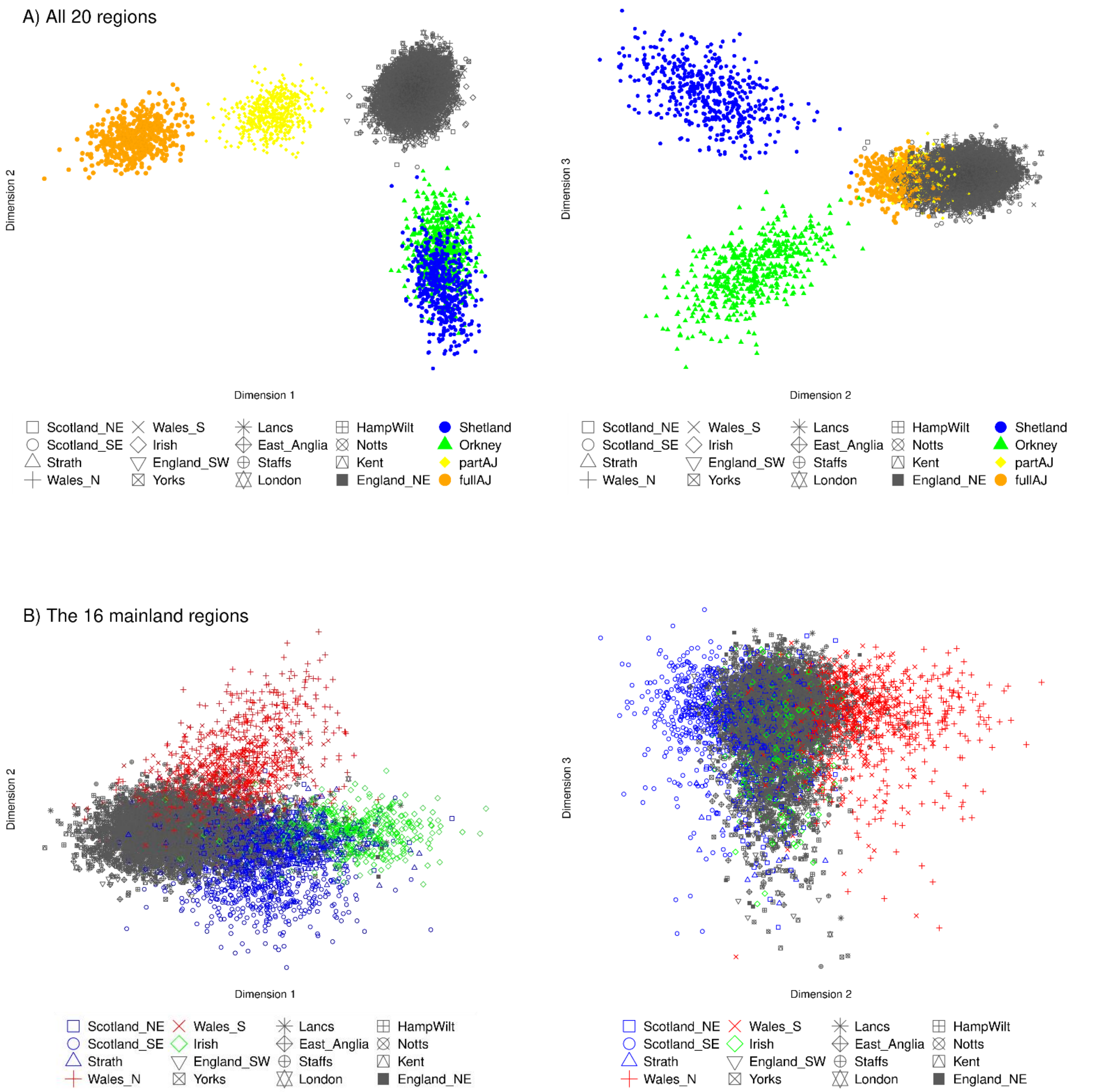


**Figure S3.** Distinction among regional populations based upon MDS analysis of rare exonic variation.

The regional MDS analyses are based on biallelic, non-singleton and LD pruned known SNPs with MAF < 5% in the considered unrelated individuals; the top 20 of the discovered MDS dimensions are subsequently used as input to the corresponding UMAP projections (Fig 1). **A)** MDS plot for all 20 groups in our study illustrating the distinction between AJ and the remaining 18 groups with part AJ group placed between full AJ and general UK population (dimension 1), the genetic distinctiveness of the Northern Isles (dimension 2) and the genetic dissimilarity between Shetlandic and Orcadian individuals (dimension 3); **B)** MDS plot for the 16 mainland regions, with dimension 1 capturing the Irish-to-Scottish/Irish-to-English difference and dimension 2 separating North Wales from South East Scotland.

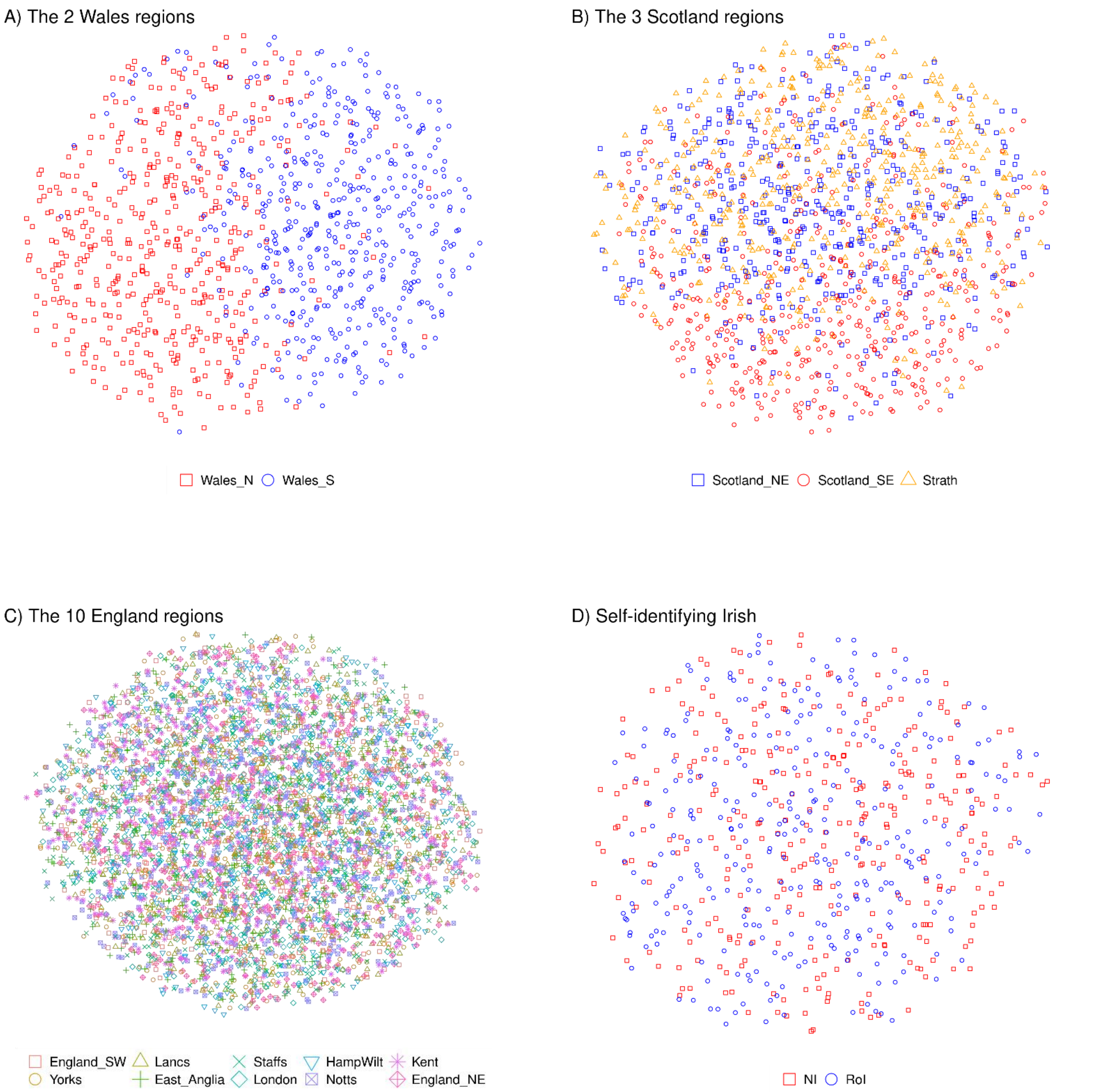


**Figure S4.** Distinction within mainland regional populations based upon UMAP projections.

The regional UMAP projections are computed on the top 20 MDS dimensions discovered based on biallelic, non-singleton and LD pruned known SNPs in unrelated individuals, with MAF thresholds adjusted according to the number of considered regions of origin. **A)** A UMAP analysis of focusing on unrelated Welsh individuals only further highlights the clear distinction between individuals born in North and South Wales (based on variants with MAF < 50%, two Welsh regions); **B)** No convincing signal for potential distinctiveness of individuals born in South East Scotland compared to the other two Scottish regions (MAF < 33%, three Scottish regions; **C)** No evident signal for genetic distinctiveness was found even for UMAP analysis of individuals born in England only (MAF < 10%, ten English regions); see also Fig S8; **D)** No evident UMAP signal (MAF < 50%, two Ireland regions) for genetic distinctiveness between self-identifying Irish individuals born in either Northern Ireland (NI, n = 333) or Republic of Ireland (RoI, n = 333)

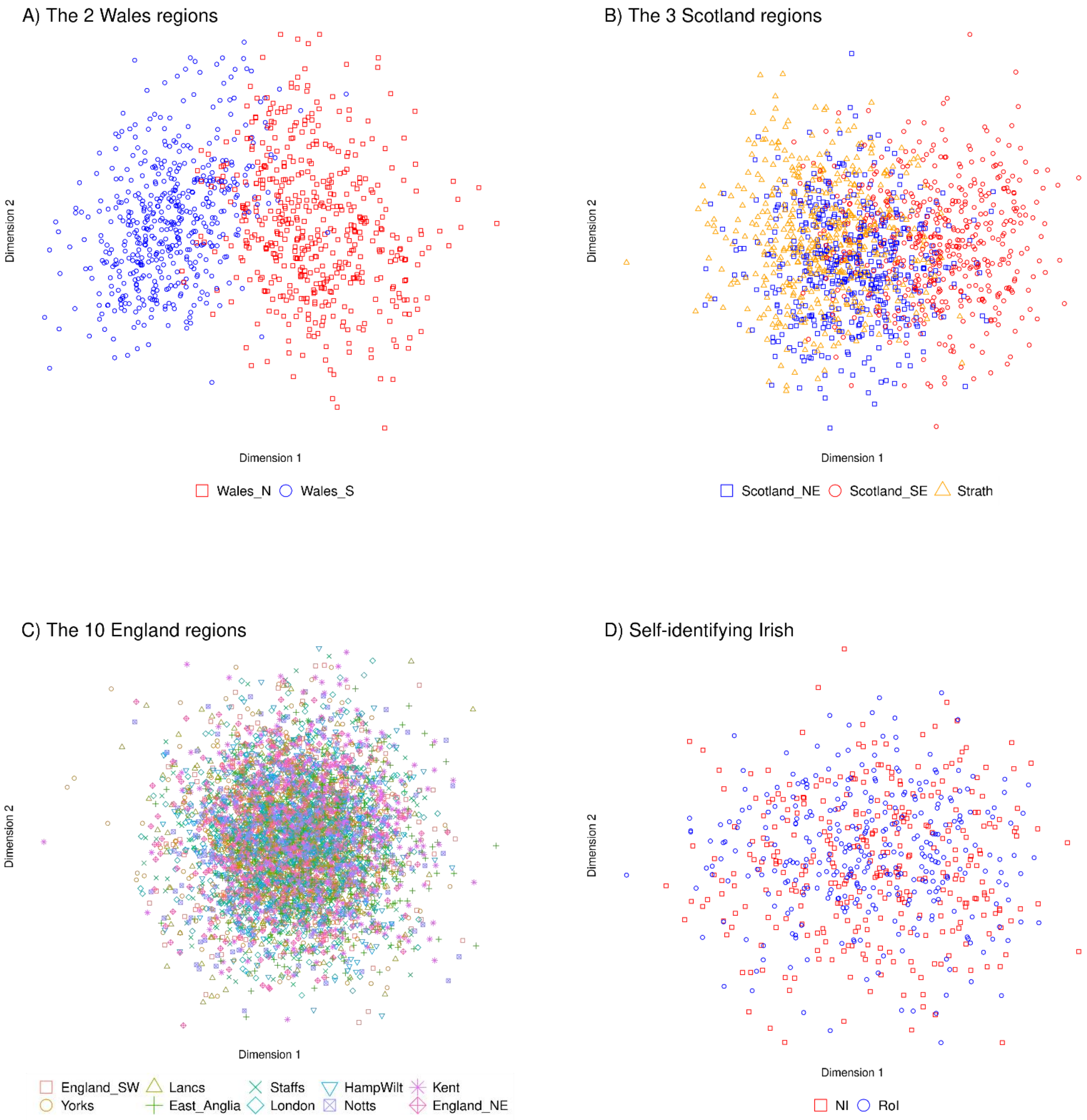


**Figure S5.** Distinction within mainland regional populations based upon MDS analysis

The regional MDS analyses are based on biallelic, non-singleton and LD pruned known SNPs in unrelated individuals, with MAF thresholds adjusted according to the number of considered regions of origin; the top 20 of the discovered MDS dimensions are subsequently used as input to the corresponding UMAP projections (Fig S4). **A)** MDS analysis focused only on Welsh individuals further confirms the North vs South split (dimension 1, based on variants with MAF < 50%, two Welsh regions); **B)** Some weak signal separating individuals born in South East Scotland from the remaining two Scottish regions is observed by focusing the MDS analysis only to individuals born in Scotland (dimension 1, MAF < 33%, three Scottish regions); **C**) no regional distinctions is observed by England-focused MDS analysis (MAF < 10%, ten English regions); **D)** no place-of-birth association is observed for self-identifying Irish individuals(MAF < 50%, two Ireland regions) born in either Northern Ireland (NI, n = 333) or Republic of Ireland (RoI, n = 333).

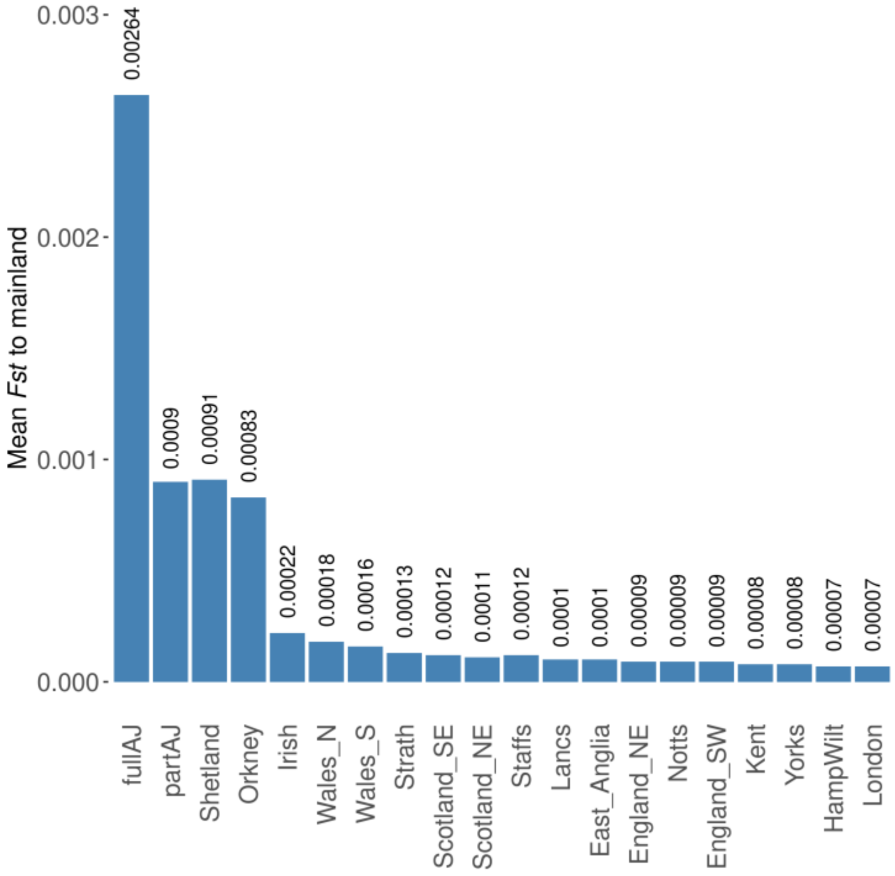


**Figure S6.** The mean $F_{ST}$ distance for each region to the 16 mainland regions (excluding the Northern Isles and AJ).

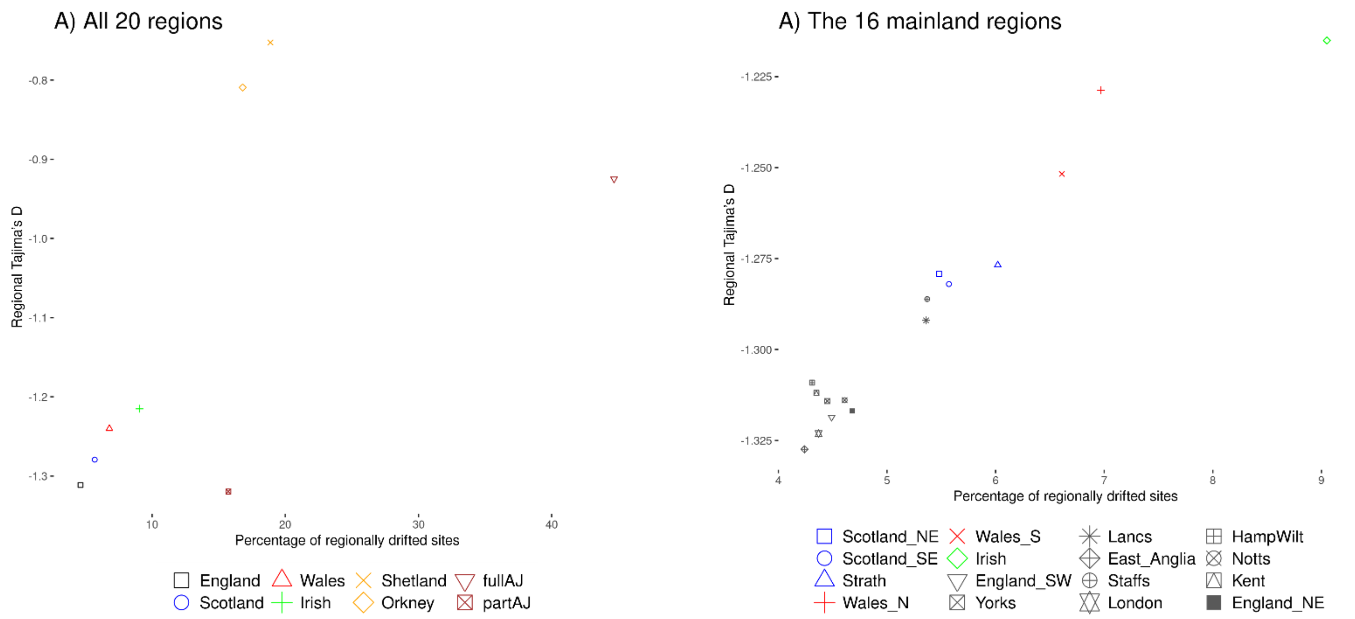


**Figure S7.** Correlation between genetic drift and Tajima's D

The x-axis represents the proportion of rare known SNPs which exhibit significant drift (at least two-fold, in both directions) in the regional cohorts compared to gnomAD NFE individuals and the y-axis correspond to the regional Tajima's D values. **A)** The analysis of all 20 regions shows a clear correlation between the genetic drift and Tajima's D measures for the mainland regions and the Northern Isles (the values for England, Scotland and Wales are computed as the average over their constitutive regions), while the unique history of full and part AJ subpopulations lead to much lower Tajima's D values than can be expected from their genetic drift measurements; **B)** Focusing on the 16 mainland regions reveals English, Scottish and Welsh regions generally cluster together, with the exception of Lancashire and Staffordshire which appear closer to the Scottish regions than they are to the remaining eight English regions. The slightly lower than expected Tajima's D values for the Irish individuals may be due to non-Irish ancestry for some of the participants in our study; in contrast to our requirement for individuals in England, Scotland and Wales to exhibit very similar genetic ancestry (i.e. "genomically British") based on a principal components analysis of the UKB whole-genome SNP array genotypes, the Irish participants were selected only based on self-identification as Irish and being born in Northern Ireland or the Republic of Ireland (Methods).

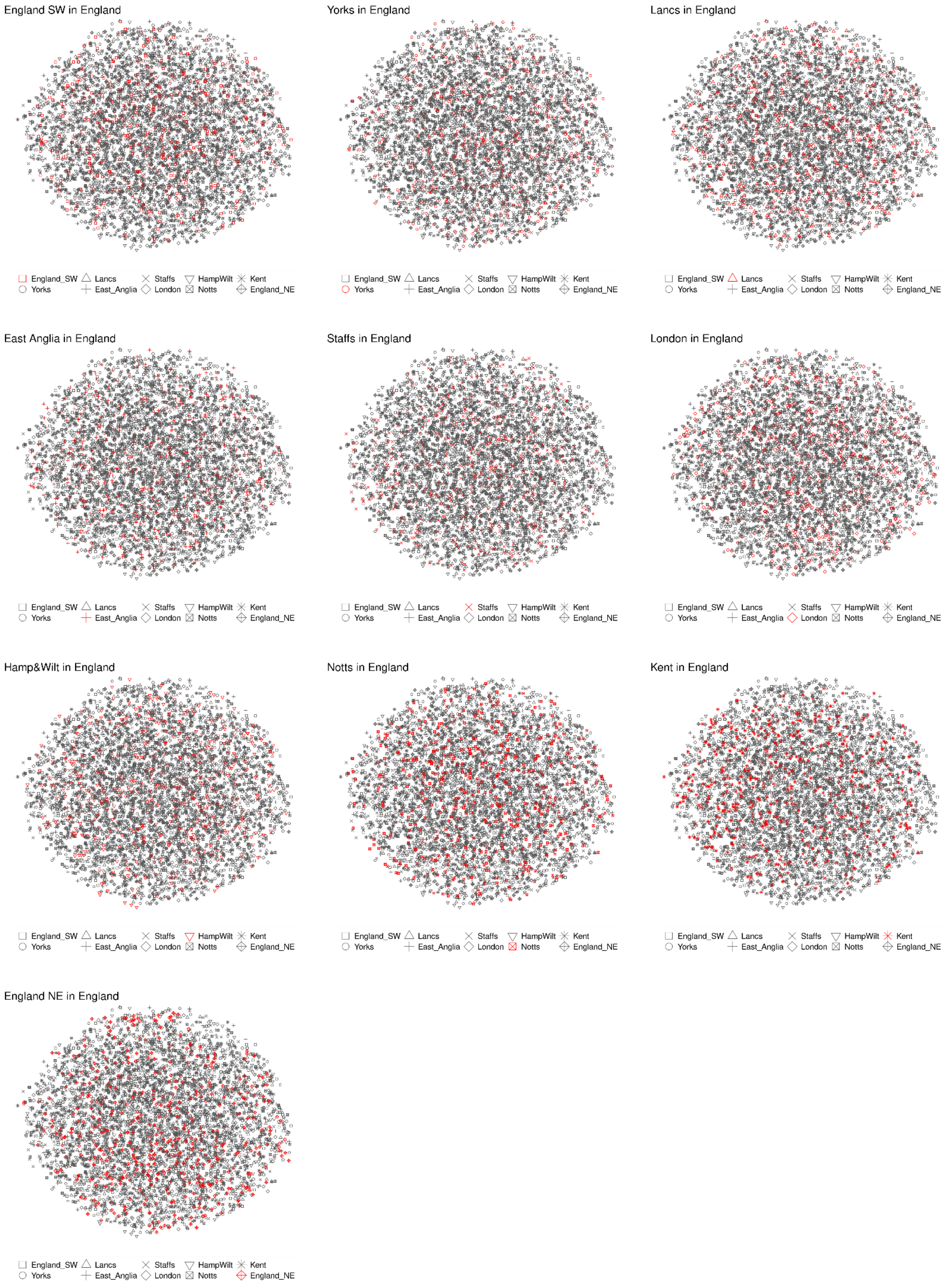


**Figure S8.** Distinction among the 10 English regions based on UMAP projections.

Same data as in Fig S4C, but coloured one region at a time for visual clarity; again no distinctive regional patterns among the 10 English regions was discovered.

### Supplementary Tables

**Table S1.** Median (IQR) counts of ALT (non-reference) alleles per individual

| | | SNP | | | | | | INDEL | | | | | |
|---|---|---|---|---|---|---|---|---|---|---|---|---|---|
| | | ultra-rare | | known | | | total | ultra-rare | | known | | | total |
| region | n | singleton | shared | very rare | rare | common | | singleton | shared | very rare | rare | common | |
| full AJ | 1004 | 11 (0) | 2 (0) | 1,037 (5) | 1,172 (5) | 30,016 (23) | 32,238 | 0 (0) | 0 (0) | 31 (1) | 34 (0) | 765 (2) | 830 |
| part AJ | 657 | 27 (1) | 1 (0) | 841 (4) | 1,146 (4) | 30,057 (23) | 32,072 | 1 (0) | 0 (0) | 27 (1) | 33 (1) | 768 (2) | 829 |
| Shetland | 492 | 16 (0) | 32 (0) | 618 (4) | 1,127 (4) | 30,071 (18) | 31,864 | 1 (0) | 2 (0) | 20 (1) | 34 (1) | 764 (2) | 821 |
| Orkney | 509 | 17 (1) | 22 (1) | 621 (3) | 1,120 (4) | 30,102 (19) | 31,882 | 1 (0) | 1 (0) | 21 (1) | 33 (1) | 764 (2) | 820 |
| Scotland NE | 1680 | 33 (1) | 6 (0) | 627 (3) | 1,119 (4) | 30,099 (21) | 31,884 | 1 (0) | 0 (0) | 20 (0) | 32 (0) | 770 (2) | 823 |
| Scotland SE | 667 | 34 (1) | 3 (1) | 627 (4) | 1,122 (4) | 30,115 (17) | 31,901 | 1 (0) | 0 (0) | 21 (1) | 32 (1) | 772 (1) | 826 |
| Strathclyde | 2077 | 29 (0) | 4 (0) | 628 (3) | 1,122 (4) | 30,099 (21) | 31,882 | 1 (0) | 0 (0) | 20 (0) | 32 (0) | 769 (2) | 822 |
| Wales N | 883 | 34 (0) | 4 (0) | 630 (3) | 1,125 (4) | 30,103 (12) | 31,896 | 1 (0) | 0 (0) | 20 (0) | 32 (1) | 770 (3) | 823 |
| Wales S | 3239 | 32 (1) | 9 (0) | 628 (2) | 1,119 (5) | 30,109 (19) | 31,897 | 1 (0) | 1 (0) | 20 (0) | 32 (0) | 773 (3) | 827 |
| Irish | 2005 | 29 (1) | 4 (0) | 618 (3) | 1,122 (5) | 30,081 (19) | 31,854 | 1 (0) | 0 (0) | 20 (0) | 32 (0) | 766 (2) | 819 |
| East Anglia | 923 | 44 (1) | 3 (0) | 634 (3) | 1,124 (4) | 30,071 (17) | 31,876 | 2 (0) | 0 (0) | 20 (0) | 33 (0) | 768 (2) | 823 |
| England NE | 2982 | 34 (0) | 8 (0) | 633 (3) | 1,124 (5) | 30,106 (21) | 31,905 | 1 (1) | 0 (0) | 21 (1) | 32 (1) | 772 (2) | 826 |
| England SW | 1412 | 41 (1) | 4 (0) | 632 (3) | 1,127 (5) | 30,091 (22) | 31,895 | 2 (0) | 0 (0) | 20 (0) | 32 (1) | 769 (2) | 823 |
| Hamp&Wilt | 1925 | 42 (0) | 3 (0) | 631 (2) | 1,124 (4) | 30,087 (20) | 31,887 | 2 (0) | 0 (0) | 20 (0) | 33 (0) | 769 (2) | 824 |
| Kent | 1327 | 43 (0) | 3 (0) | 633 (3) | 1,124 (5) | 30,082 (17) | 31,885 | 2 (0) | 0 (0) | 20 (0) | 33 (1) | 768 (2) | 823 |
| Lancs | 3007 | 31 (1) | 7 (1) | 634 (3) | 1,123 (4) | 30,065 (16) | 31,860 | 1 (0) | 1 (1) | 20 (1) | 32 (1) | 765 (3) | 819 |
| Notts | 4192 | 34 (0) | 11 (0) | 635 (3) | 1,121 (4) | 30,077 (22) | 31,878 | 1 (1) | 1 (0) | 20 (1) | 32 (1) | 768 (2) | 822 |
| Staffs | 3526 | 27 (1) | 15 (0) | 636 (2) | 1,124 (4) | 30,102 (22) | 31,904 | 1 (0) | 1 (0) | 20 (0) | 32 (1) | 771 (1) | 825 |
| Yorks | 3276 | 36 (1) | 7 (0) | 633 (3) | 1,122 (5) | 30,100 (16) | 31,898 | 2 (0) | 0 (0) | 20 (1) | 32 (1) | 770 (2) | 824 |
| London | 8913 | 36 (1) | 10 (0) | 633 (2) | 1,123 (4) | 30,067 (21) | 31,869 | 2 (1) | 1 (0) | 20 (0) | 32 (0) | 764 (2) | 819 |

**n**: number of unrelated individuals per region; **ultra-rare** SNP/INDEL: variants not found in the gnomAD dataset (v3.1.1, containing data for 76156 genomes from unrelated individuals world-wide), singleton: variant found only in a single individual from this region, shared: variant found in more than one individual from this region; **very rare** SNP/INDEL: found in gnomADg v3.1.1 dataset with Non-Finnish European (NFE, n = 34,029) MAF < 1%; **rare** SNP/INDEL: 1% ≤ NFE MAF < 5%; **common** SNP/INDEL: NFE MAF ≥ 5%; **median and IQR** values for each region: based on random selection of 450 individuals (with replacement) repeated 10k times

**Table S2.** Pair-wise $F_{ST}$ estimates for the 20 UKB regions

| | full AJ | part AJ | Shetland | Orkney | Irish | Wales N | Wales S | Strathclyde | Scotland SE | Scotland NE | Staffs | Lancs | East Anglia | England NE | Notts | England SW | Kent | Yorks | Hamp&Wilt | London |
|---|---|---|---|---|---|---|---|---|---|---|---|---|---|---|---|---|---|---|---|---|
| full AJ | n/a | 0.00068 | 0.00382 | 0.00371 | 0.00288 | 0.00277 | 0.00275 | 0.00267 | 0.00266 | 0.00265 | 0.00263 | 0.00262 | 0.00256 | 0.00258 | 0.00257 | 0.00260 | 0.00258 | 0.00257 | 0.00257 | 0.00256 |
| part AJ | 0.00068 | n/a | 0.00172 | 0.00165 | 0.00107 | 0.00100 | 0.00097 | 0.00094 | 0.00093 | 0.00092 | 0.00090 | 0.00089 | 0.00084 | 0.00087 | 0.00085 | 0.00086 | 0.00085 | 0.00085 | 0.00084 | 0.00083 |
| Shetland | 0.00382 | 0.00172 | n/a | 0.00132 | 0.00101 | 0.00102 | 0.00101 | 0.00087 | 0.00087 | 0.00084 | 0.00094 | 0.00091 | 0.00090 | 0.00088 | 0.00090 | 0.00089 | 0.00089 | 0.00086 | 0.00087 | 0.00087 |
| Orkney | 0.00371 | 0.00165 | 0.00132 | n/a | 0.00091 | 0.00094 | 0.00092 | 0.00078 | 0.00078 | 0.00075 | 0.00087 | 0.00083 | 0.00082 | 0.00080 | 0.00082 | 0.00082 | 0.00081 | 0.00079 | 0.00079 | 0.00079 |
| Irish | 0.00288 | 0.00107 | 0.00101 | 0.00091 | n/a | 0.00029 | 0.00028 | 0.00012 | 0.00017 | 0.00015 | 0.00029 | 0.00024 | 0.00026 | 0.00020 | 0.00026 | 0.00023 | 0.00024 | 0.00021 | 0.00022 | 0.00020 |
| Wales N | 0.00277 | 0.00100 | 0.00102 | 0.00094 | 0.00029 | n/a | 0.00015 | 0.00022 | 0.00022 | 0.00020 | 0.00017 | 0.00017 | 0.00018 | 0.00017 | 0.00017 | 0.00017 | 0.00017 | 0.00016 | 0.00015 | 0.00014 |
| Wales S | 0.00275 | 0.00097 | 0.00101 | 0.00092 | 0.00028 | 0.00015 | n/a | 0.00020 | 0.00021 | 0.00019 | 0.00018 | 0.00017 | 0.00015 | 0.00015 | 0.00015 | 0.00013 | 0.00013 | 0.00014 | 0.00012 | 0.00012 |
| Strathclyde | 0.00267 | 0.00094 | 0.00087 | 0.00078 | 0.00012 | 0.00022 | 0.00020 | n/a | 0.00004 | 0.00004 | 0.00017 | 0.00014 | 0.00014 | 0.00009 | 0.00013 | 0.00013 | 0.00013 | 0.00010 | 0.00011 | 0.00011 |
| Scotland SE | 0.00266 | 0.00093 | 0.00087 | 0.00078 | 0.00017 | 0.00022 | 0.00021 | 0.00004 | n/a | 0.00004 | 0.00016 | 0.00014 | 0.00013 | 0.00008 | 0.00012 | 0.00011 | 0.00011 | 0.00009 | 0.00010 | 0.00010 |
| Scotland NE | 0.00265 | 0.00092 | 0.00084 | 0.00075 | 0.00015 | 0.00020 | 0.00019 | 0.00004 | 0.00004 | n/a | 0.00016 | 0.00012 | 0.00012 | 0.00008 | 0.00011 | 0.00010 | 0.00010 | 0.00008 | 0.00009 | 0.00008 |
| Staffs | 0.00263 | 0.00090 | 0.00094 | 0.00087 | 0.00029 | 0.00017 | 0.00018 | 0.00017 | 0.00016 | 0.00016 | n/a | 0.00008 | 0.00009 | 0.00010 | 0.00006 | 0.00009 | 0.00008 | 0.00008 | 0.00007 | 0.00007 |
| Lancs | 0.00262 | 0.00089 | 0.00091 | 0.00083 | 0.00024 | 0.00017 | 0.00017 | 0.00014 | 0.00014 | 0.00012 | 0.00008 | n/a | 0.00007 | 0.00007 | 0.00005 | 0.00008 | 0.00006 | 0.00005 | 0.00006 | 0.00006 |
| East Anglia | 0.00256 | 0.00084 | 0.00090 | 0.00082 | 0.00026 | 0.00018 | 0.00015 | 0.00014 | 0.00013 | 0.00012 | 0.00009 | 0.00007 | n/a | 0.00007 | 0.00004 | 0.00005 | 0.00004 | 0.00004 | 0.00003 | 0.00002 |
| England NE | 0.00258 | 0.00087 | 0.00088 | 0.00080 | 0.00020 | 0.00017 | 0.00015 | 0.00009 | 0.00008 | 0.00008 | 0.00010 | 0.00007 | 0.00007 | n/a | 0.00005 | 0.00006 | 0.00005 | 0.00003 | 0.00005 | 0.00004 |
| Notts | 0.00257 | 0.00085 | 0.00090 | 0.00082 | 0.00026 | 0.00017 | 0.00015 | 0.00013 | 0.00012 | 0.00011 | 0.00006 | 0.00005 | 0.00004 | 0.00005 | n/a | 0.00005 | 0.00003 | 0.00003 | 0.00003 | 0.00002 |
| England SW | 0.00260 | 0.00086 | 0.00089 | 0.00082 | 0.00023 | 0.00017 | 0.00013 | 0.00013 | 0.00011 | 0.00010 | 0.00009 | 0.00008 | 0.00005 | 0.00006 | 0.00005 | n/a | 0.00004 | 0.00005 | 0.00002 | 0.00002 |
| Kent | 0.00258 | 0.00085 | 0.00089 | 0.00081 | 0.00024 | 0.00017 | 0.00013 | 0.00013 | 0.00011 | 0.00010 | 0.00008 | 0.00006 | 0.00004 | 0.00005 | 0.00003 | 0.00004 | n/a | 0.00004 | 0.00002 | 0.00002 |
| Yorks | 0.00257 | 0.00085 | 0.00086 | 0.00079 | 0.00021 | 0.00016 | 0.00014 | 0.00010 | 0.00009 | 0.00008 | 0.00008 | 0.00005 | 0.00004 | 0.00003 | 0.00003 | 0.00005 | 0.00004 | n/a | 0.00003 | 0.00003 |
| Hamp&Wilt | 0.00257 | 0.00084 | 0.00087 | 0.00079 | 0.00022 | 0.00015 | 0.00012 | 0.00011 | 0.00010 | 0.00009 | 0.00007 | 0.00006 | 0.00003 | 0.00005 | 0.00003 | 0.00002 | 0.00002 | 0.00003 | n/a | 0.00001 |
| London | 0.00256 | 0.00083 | 0.00087 | 0.00079 | 0.00020 | 0.00014 | 0.00012 | 0.00011 | 0.00010 | 0.00008 | 0.00007 | 0.00006 | 0.00002 | 0.00004 | 0.00002 | 0.00002 | 0.00002 | 0.00003 | 0.00001 | n/a |

| $F_{ST}$ values |
|---|
| < 0.0001 |
| 0.0001-0.0002 |
| 0.0002-0.0003 |
| 0.0003 - 0.001 |
| > 0.001 |

**Table S3.** Genetic drift observed in the 20 UKB regions w.r.t. Non-Finnish Europeans (NFE) in gnomADg 3.1.1. (sorted by total proportion of up-drifted sites)

| | **down-drift** | | **no change** | **up-drift** | | |
|---|---|---|---|---|---|---|
| | **> 4 fold** | **2-4 fold** | | **2-4 fold** | **> 4 fold** | **total up-drift** |
| full AJ | 14.68% | 17.32% | 55.31% | 11.19% | 1.50% | 12.69% |
| part AJ | 2.00% | 8.90% | 84.28% | 4.65% | 0.17% | 4.82% |
| Shetland | 3.62% | 11.64% | 81.12% | 3.53% | 0.09% | 3.62% |
| Orkney | 3.39% | 10.35% | 83.19% | 2.98% | 0.08% | 3.06% |
| Irish | 1.72% | 6.15% | 90.94% | 1.10% | 0.08% | 1.18% |
| Wales N | 1.40% | 4.80% | 93.03% | 0.70% | 0.07% | 0.77% |
| Wales S | 1.39% | 4.74% | 93.39% | 0.41% | 0.07% | 0.48% |
| Scotland SE | 1.29% | 3.85% | 94.44% | 0.36% | 0.07% | 0.43% |
| Strathclyde | 1.40% | 4.21% | 93.98% | 0.34% | 0.07% | 0.41% |
| Scotland NE | 1.32% | 3.79% | 94.51% | 0.31% | 0.06% | 0.37% |
| Lancs | 1.27% | 3.74% | 94.65% | 0.28% | 0.07% | 0.35% |
| Staffs | 1.26% | 3.77% | 94.64% | 0.27% | 0.07% | 0.34% |
| England NE | 1.18% | 3.20% | 95.32% | 0.23% | 0.07% | 0.30% |
| Notts | 1.24% | 3.09% | 95.39% | 0.23% | 0.05% | 0.28% |
| Yorks | 1.10% | 3.09% | 95.55% | 0.19% | 0.07% | 0.26% |
| Kent | 1.26% | 2.86% | 95.65% | 0.16% | 0.07% | 0.23% |
| England SW | 1.15% | 3.13% | 95.52% | 0.14% | 0.07% | 0.21% |
| Hamp&Wilt | 1.27% | 2.83% | 95.69% | 0.13% | 0.08% | 0.21% |
| London | 1.18% | 2.99% | 95.62% | 0.14% | 0.06% | 0.20% |
| East Anglia | 1.16% | 2.89% | 95.75% | 0.13% | 0.06% | 0.19% |

**Table S4.** Tajima's D median and first/third quartile values observed in the 20 UKB regions (sorted by median)

| **region** | **median** | **25%** | **75%** |
|---|---|---|---|
| Shetland | -0.75265 | -1.17 | -0.31 |
| Orkney | -0.80941 | -1.21 | -0.39 |
| full AJ | -0.92492 | -1.31 | -0.54 |
| Irish | -1.21509 | -1.54 | -0.84 |
| Wales N | -1.22871 | -1.57 | -0.87 |
| Wales S | -1.25174 | -1.58 | -0.88 |
| Strathclyde | -1.27679 | -1.60 | -0.90 |
| Scotland NE | -1.27920 | -1.61 | -0.89 |
| Scotland SE | -1.28201 | -1.59 | -0.91 |
| Staffs | -1.28613 | -1.60 | -0.91 |
| Lancs | -1.29199 | -1.61 | -0.94 |
| Hamp&Wilt | -1.30909 | -1.64 | -0.95 |
| Kent | -1.31190 | -1.63 | -0.95 |
| Notts | -1.31390 | -1.63 | -0.95 |
| Yorks | -1.31415 | -1.64 | -0.95 |
| England NE | -1.31687 | -1.63 | -0.94 |
| England SW | -1.31859 | -1.63 | -0.96 |
| part AJ | -1.31949 | -1.63 | -0.96 |
| London | -1.32309 | -1.65 | -0.98 |
| East Anglia | -1.32740 | -1.63 | -0.97 |

**Table S5.** Variant QC filtering statistics, reporting the number of sites filtered at each step (as a proportion of the sites submitted to it)

| | | Filtering based on | | | | Filtering based on DP, GQ and VAF | | | |
|---|---|---|---|---|---|---|---|---|---|
| **Region** | **miss≥10%** | **not in target regions** | **gnomAD FAIL** | **in LCR** | **SNPs in PCW** | **individual SNP calls** | **individual INDEL calls** | **SNP sites** | **INDEL sites** |
| Shetland | 4.35% | 54.99% | 3.94% | 1.37% | 0.48% | 0.78% | 4.51% | 0.50% | 1.92% |
| Orkney | 4.16% | 54.89% | 3.85% | 1.38% | 0.48% | 0.78% | 4.59% | 0.47% | 1.77% |
| Scotland SE | 3.38% | 54.06% | 3.30% | 1.29% | 0.42% | 0.70% | 4.00% | 0.74% | 2.85% |
| Scotland NE | 2.58% | 52.87% | 2.73% | 1.16% | 0.39% | 0.71% | 4.17% | 0.70% | 2.82% |
| Strathclyde | 2.38% | 52.62% | 2.60% | 1.15% | 0.39% | 0.72% | 4.10% | 0.68% | 2.97% |
| Wales N | 3.33% | 53.75% | 3.19% | 1.26% | 0.42% | 0.73% | 4.37% | 0.77% | 2.98% |
| Wales S | 2.04% | 52.10% | 2.35% | 1.06% | 0.37% | 0.70% | 4.10% | 0.67% | 2.66% |
| Irish | 2.63% | 52.76% | 2.72% | 1.18% | 0.39% | 0.74% | 4.37% | 0.77% | 3.22% |
| East Anglia | 3.05% | 53.47% | 3.01% | 1.18% | 0.39% | 0.75% | 4.39% | 0.74% | 3.20% |
| England NE | 1.97% | 52.14% | 2.31% | 1.07% | 0.35% | 0.71% | 4.13% | 0.66% | 2.77% |
| England SW | 2.61% | 52.88% | 2.71% | 1.16% | 0.39% | 0.73% | 4.25% | 0.69% | 3.00% |
| Hamp&Wilt | 2.28% | 52.48% | 2.49% | 1.13% | 0.37% | 0.72% | 4.25% | 0.71% | 3.03% |
| Kent | 2.68% | 52.98% | 2.72% | 1.15% | 0.38% | 0.75% | 4.26% | 0.71% | 3.29% |
| Lancs | 2.12% | 52.20% | 2.40% | 1.08% | 0.36% | 0.75% | 4.39% | 0.70% | 2.87% |
| Yorks | 1.88% | 51.98% | 2.24% | 1.06% | 0.35% | 0.71% | 4.18% | 0.63% | 2.66% |
| Notts | 1.77% | 51.78% | 2.13% | 1.01% | 0.34% | 0.73% | 4.30% | 0.66% | 2.79% |
| Staffs | 2.00% | 52.20% | 2.35% | 1.07% | 0.38% | 0.71% | 4.15% | 0.65% | 2.92% |
| London | 1.37% | 51.05% | 1.77% | 0.89% | 0.29% | 0.75% | 4.41% | 0.58% | 2.26% |
| all AJ | 3.08% | 54.03% | 3.06% | 1.21% | 0.45% | 0.75% | 4.46% | 0.78% | 3.31% |

**miss≥10%**: sites excluded due to more than 10% of the individuals are with missing genotype

**not in target regions**: sites excluded due to variant location being outside of the WES capture target region

**gnomAD FAIL**: sites excluded due to being found to fail the gnomADg QC

**in LCR**: sites excluded due to variant location in Low-Complexity Region (LCR)

**SNPs in PCW**: SNP sites excluded due to being in Poor Coverage Window (PCW) region in gnomADg (defined as 10bp windows centred on any base with coverage < 10x)

**individual SNP calls**: individual SNP genotypes being reset to REF due to poor DP, GQ and/or VAF; leading to excluding the corresponding number of **SNP sites**

**individual INDEL calls**: individual INDEL genotypes being reset to REF due to poor DP, GQ and/or VAF; leading to excluding the corresponding number of **INDEL sites**
